# Dysregulated Wnt and NFAT signaling in a Parkinson’s disease LRRK2 G2019S knock-in model

**DOI:** 10.1101/2023.03.31.535090

**Authors:** Andrea Wetzel, Si Hang Lei, Tiansheng Liu, Michael P. Hughes, Yunan Peng, Tristan McKay, Simon N. Waddington, Simone Grannò, Ahad A. Rahim, Kirsten Harvey

## Abstract

**Background:** Parkinson’s disease (PD) is a progressive late-onset neurodegenerative disease leading to physical and cognitive decline. Mutations of leucine-rich repeat kinase 2 (*LRRK2*) are the most common genetic cause of PD. LRRK2 is a complex scaffolding protein with known regulatory roles in multiple molecular pathways. Two prominent examples of LRRK2-modulated pathways are Wingless/Int (Wnt) and nuclear factor of activated T-cells (NFAT) signaling. Both are well described key regulators of immune and nervous system development as well as maturation. The aim of this study was to establish the physiological and pathogenic role of LRRK2 in Wnt and NFAT signaling in the brain, as well as the potential contribution of the non-canonical Wnt/Calcium pathway.

**Methods:** *In vivo* cerebral Wnt and NFATc1 signaling activity was quantified in LRRK2 G2019S mutant knock-in (KI) and LRRK2 knockout (KO) male and female mice with repeated measures over 28 weeks, employing lentiviral luciferase biosensors, and analyzed using a mixed-effect model. To establish spatial resolution, we investigated tissues, and primary neuronal cell cultures from different brain regions combining luciferase signaling activity, immunohistochemistry, qPCR and western blot assays. Results were analyzed by unpaired t-test with Welch’s correction or 2-way ANOVA with post hoc corrections.

**Results:** *In vivo* Wnt signaling activity in LRRK2 KO and LRRK2 G2019S KI mice was increased significantly ∼3-fold, with a more pronounced effect in males (∼4-fold) than females (∼2-fold). NFATc1 signaling was reduced ∼0.5-fold in LRRK2 G2019S KI mice. Brain tissue analysis showed region-specific expression changes in Wnt and NFAT signaling components. These effects were predominantly observed at the protein level in the striatum and cerebral cortex of LRRK2 KI mice. Primary neuronal cell culture analysis showed significant genotype-dependent alterations in Wnt and NFATc1 signaling under basal and stimulated conditions. Wnt and NFATc1 signaling was primarily dysregulated in cortical and hippocampal neurons respectively.

**Conclusions:** Our study further built on knowledge of LRRK2 as a Wnt and NFAT signaling protein. We identified complex changes in neuronal models of LRRK2 PD, suggesting a role for mutant LRRK2 in the dysregulation of NFAT, and canonical and non-canonical Wnt signaling.

## Background

Parkinson’s disease (PD) is the second most common neurodegenerative disease worldwide. In light of the current trend of ageing populations, the healthcare burden of PD is projected to increase substantially over the next decades. Whilst available medications have improved prognosis over recent years, definitive treatment remains unavailable. These notions underscore the urgent need to identify novel therapeutic targets.

Whilst the onset of PD is mostly sporadic, 10% of PD cases display genetic inheritance (1). Mutations in the Leucine-rich repeat kinase 2 gene (*LRRK2)* are the most common cause of familial PD and genome-wide association studies revealed that the *LRRK2* locus contains also risk factors for sporadic PD (2)(3)(4). LRRK2 is a large enzyme with kinase and GTPase activities, and is an extensive binding partner for more than 260 proteins, involved in multiple cellular processes and signaling pathways (5)(6)(7). Even though LRRK2 has been investigated for nearly two decades, its ultimate function has not been clarified entirely. Efforts towards understanding LRRK2 pathophysiology have gained traction since its link to PD was first established in 2004 (8)(9).

The most prevalent *LRRK2* mutation leads to the LRRK2 G2019S variant in exon 41, occurring in 2-6% and 1-2% of familial and sporadic cases, respectively (10). The encoded mutant protein reportedly displays enhanced LRRK2 kinase activity (11)(12)(13)(14), which led to the development of LRRK2 kinase inhibitors as potential therapeutic approach.

Binding to a large number of cellular partners, LRRK2 has been shown to act as a scaffolding protein, facilitating the interplay between diverse signaling mediators (14)(15)(16)(17). Amongst pathways suggested to be regulated by LRRK2 are the evolutionarily highly conserved Wingless/Integrated (Wnt) signaling cascades (18)(19) as well as the nuclear factor of activated T-cells (NFAT) pathway (14)(20)(21). Both share several key molecules such as glycogen synthase kinase 3β (GSK-3β) and casein kinase 1 (CK1), and are critical for immune and nervous system development, adult homeostasis and tissue regeneration (17)(22)(23)(24)(25). Wnt signaling is generally distinguishable as (A) the canonical Wnt pathway, also known as Wnt/β-catenin pathway, (B) the Wnt/planar cell polarity (PCP) pathway, and (C) the Wnt/Calcium pathway. LRRK2 has been reported to act as scaffold in all three major branches of Wnt signaling by binding to cytoplasmic disheveled (DVL) proteins, which are important throughout pathway activation (6)(14)(18)(26)(27). To activate Wnt signaling, Wnt ligands bind to membrane bound Frizzled (Fz) receptors, triggering a conformational change resulting in recruitment and partial phosphorylation of pathway-specific intracellular signaling molecules, including DVLs. Activation of the canonical Wnt pathway commonly requires the membrane-bound Wnt co-receptor low-density lipoprotein receptor-related protein 5/6 (LRP5/6) (28). While Wnt3a and Wnt7a primarily stimulate canonical Wnt signaling activity, Wnt5a promotes non-canonical Wnt signaling activity (27)(29). Moreover, Wnt5a has previously been shown to also inhibit canonical Wnt signaling via distinct and specific receptors (30)(31).

Previous investigations have suggested LRRK2 to support protein complex formation after Wnt pathway activation as signaling scaffold by binding to DVLs, LRP6 and components of the β-catenin destruction complex (BDC), such as Axin1, β-catenin and GSK-3β (18). As a result of canonical Wnt signaling activation, free β-catenin moves to the nucleus inducing gene transcription by binding to nuclear transcription factors such as T-cell specific transcription factor/lymphoid enhancer binding factor (TCF/LEF) (32).

In the case of NFAT signaling, GSK-3β and LRRK2 have been reported to be part of an inhibitory complex called NRON (21). LRRK2 again acts as scaffolding protein bridging and stabilizing this NFAT-NRON complex protein-protein interaction. Interestingly, Wnt ligand binding to Frizzled and co-receptors can also activate NFAT signaling via an increase of cytoplansmic Ca^2+^.

Dysregulation of Wnt and NFAT pathways, as well as genetic changes in *LRRK2* are associated with several pathological processes including cancer, infectious, immunological and neurodegenerative disorders (32)(33)(34)(35)(36)(37)(38)(39)(40)(41)(42)(43)(44)(45). Understanding the physiological and pathogenic role of wild type and mutant LRRK2 as a regulator of Wnt and NFAT signaling would therefore broaden our knowledge and support the identification of novel therapeutic strategies.

In this study, we investigated the role of LRRK2 in Wnt and NFAT signaling *in vivo* by monitoring both pathways in LRRK2 knockout and LRRK2 G2019S knock-in mice by using a lentiviral construct that allowed visualization of the transcriptional activity of TCF/LEF and NFAT. We then showed that several brain regions feature LRRK2-dependent Wnt and NFAT signaling dysregulation at the transcriptional and protein level. Finally, we assessed cell type-specific effects of Wnt and NFAT stimulation in different LRRK2 genotypes in primary murine neuronal cell cultures. Throughout this study, we also segregated the data by sex, in order to better elucidate potential sex differences.

## Methods

### Animals

All animal procedures in this study were performed upon approval by the University College London Animal Welfare and Ethical Review Body and licensed by the UK Home Office (PPL 80/2486 and PPL 70/9070). LRRK2 knockout mice (B6.129X1 (FVB)-Lrrk2^tm1.1Cai/J^) from the Jackson Laboratory and LRRK2 G2019S knock-in mice, kindly provided by Heather L. Melrose, were bred and maintained at the UCL School of Pharmacy. Exon 2 of the *Lrrk2* gene has been deleted in LRRK2 knockout mice, resulting in a premature stop codon in exon 3 (46). Provided protocols from the Jackson Laboratory were used for *Lrrk2* knockout mouse genotyping. LRRK2 G2019S knock-in mice were generated by replacing two bases in exon 41 of the *Lrrk2* gene (47). Genotyping of LRRK2 G2019S knock-in mice was performed using primers that detect the specific knock-in mutation and the remaining 34 base loxP sequence (47). Both mouse lines were bred against a C57BL/6J background.

### Lentiviral reporter constructs and lentivirus production

Transcription factor activated reporter lentiviral constructs were used to investigate Wnt and NFATc1 signaling activity *in vitro* and *in vivo*. Biosensor construct generation has been described in detail previously (48). The lentiviral construct is based on a second generation cassette containing U3 deleted 3’-long terminal repeats (LTRs). In between the LTRs exists a gateway cloning site upstream of a bicistronic *in vivo* optimized firefly luciferase-2A-e green fluorescent protein (GFP) reporter cassette. An upstream central polypurine tract (cPPT) and a downstream woodchuck hepatitis virus post-transcriptional regulatory element (WPRE) enhance gene expression. Serial transcription factor binding sequences for the TCF/LEF (8x AGATCAAAGGGGGTA) and NFATc1 transcription factor (4x GGAGGAAAAACTGTTTCATACAGAAGGCGT) were cloned upstream of the minimal promoter driven luciferase in the gateway region (pLNT-TCF/LEF or NFAT-FLuc-2A-eGFP-JDG) (48). The lentiviral expression construct with a SFFV constitutive promoter, driving expression of the firefly luciferase-2A-eGFP bicistronic reporter (pLNT-SFFV-FLuc-2A-eGFP-JDG), served as positive control in our *in vivo* experiments. For our *in vitro* analysis, we used a TCF/LEF and NFATc1 activated biosensor construct that induces the expression of a secreted Nanoluciferase-2A-eGFP bicistronic reporter (pLNT-TCF/LEF or NFAT-secNanoLuc-2A-GFP). As positive and loading control served a lentiviral construct with the SFFV constructive promoter, driving the expression of a secreted vargulin luciferase (pLNT-SFFV-secVLuc). Different substrates are required for the corresponding construct.

For lentivirus production, HEK293T cells were seeded overnight in T175 cm^2^ flasks and triple transfected with 50µg transcription factor activated reporter plasmid, 17.5µg VSV-G envelope plasmid (pMD.G2), and 32.5µg gag-pol packaging plasmid (pCMVΔR8.74) using 10 mM polyethylenimine (Sigma) in OptiMEM for three hours (48). Media was changed to DMEM with 10% fetal bovine serum and lentiviral supernatants were harvested 48 and 72 hours after transfection. The supernatant was filtered using 0.22µm sterile filters (Millipore) and lentiviral particles were concentrated by low-g centrifugation overnight. Suspended vector was stored at −80°C and vector titre was determined with the p24 assay from Zeptometrix.

### Animal procedures and bioluminescence imaging of mice

LRRK2 knockout, LRRK2 G2019S knock-in and wild type mice were time mated to generate neonatal pups. Neonates were momentarily anesthetized on ice at postnatal day one and injected with 5µl of a concentrated lentivirus suspension (1×10^9^ vector particles/ml) via intracerebroventricular (ICV) injection targeting the anterior horn of the lateral ventricle by using a 33-gauge needle (Hamilton, Reno, NV, USA) (49)(50). The lentivirus contained either the pLNT-TCF/LEF-FLuc-2A-eGFP-JDG or pLNT-NFAT-FLuc-2A-eGFP-JDG construct coding for the transcription factor activated firefly luciferase and GFP (or pLNT-SFFV-FLuc-2A-eGFP-JDG as positive control). Injected postnatal day one old mice were returned to the dam and after one week injected intraperitoneally with 150mg/kg firefly D-luciferin (sterile filtered 15mg/ml in phosphate buffered saline, Gold Biotechnology) (48) and imaged 10 minutes after D-luciferin injection with a cooled charge-coupled device (CCD) camera (In Vivo Imaging System by PerkinElmer). Mice were imaged regularly every week until week 8 and every 4 weeks until week 28. Grey-scale photographs were taken followed by bioluminescence images using a suitable, for all genotypes consistent binning (resolution) factor per age with a 1.2/f stop. The regions of interest (ROIs) were defined manually and signal intensities were calculated with Living Image software (Perkin Elmer) and expressed as photons per seconds (48). To identify an optimal time frame for luciferase activity measurements, injected mice were anesthetized with isoflurane (Zoetis UK Limited) and bioluminescence images were taken every two minutes for a period of 52 minutes, resulting in an optimal measurement time frame of 10 to 20 minutes after D-luciferin injection. This is the time frame when luciferin has reached maximum biodistribution (Figure 1D). Non injected littermates of each genotype served as negative control throughout the study. After 6 months, mice were injected with D-luciferin via intraperitoneal injection, sacrificed and individual organs (brain, heart, kidney, stomach, lung, spleen) were extracted and imaged with the IVIS machine (Figure 1C). Shortly after, organs were snap frozen on dry ice and stored at −80°C for further vector copy number analysis. Alternatively, vector injected and non-injected mice were perfused with phosphate buffered saline (PBS) and fixed in 4% paraformaldehyde for immunohistochemistry analysis.

**Figure 1.**
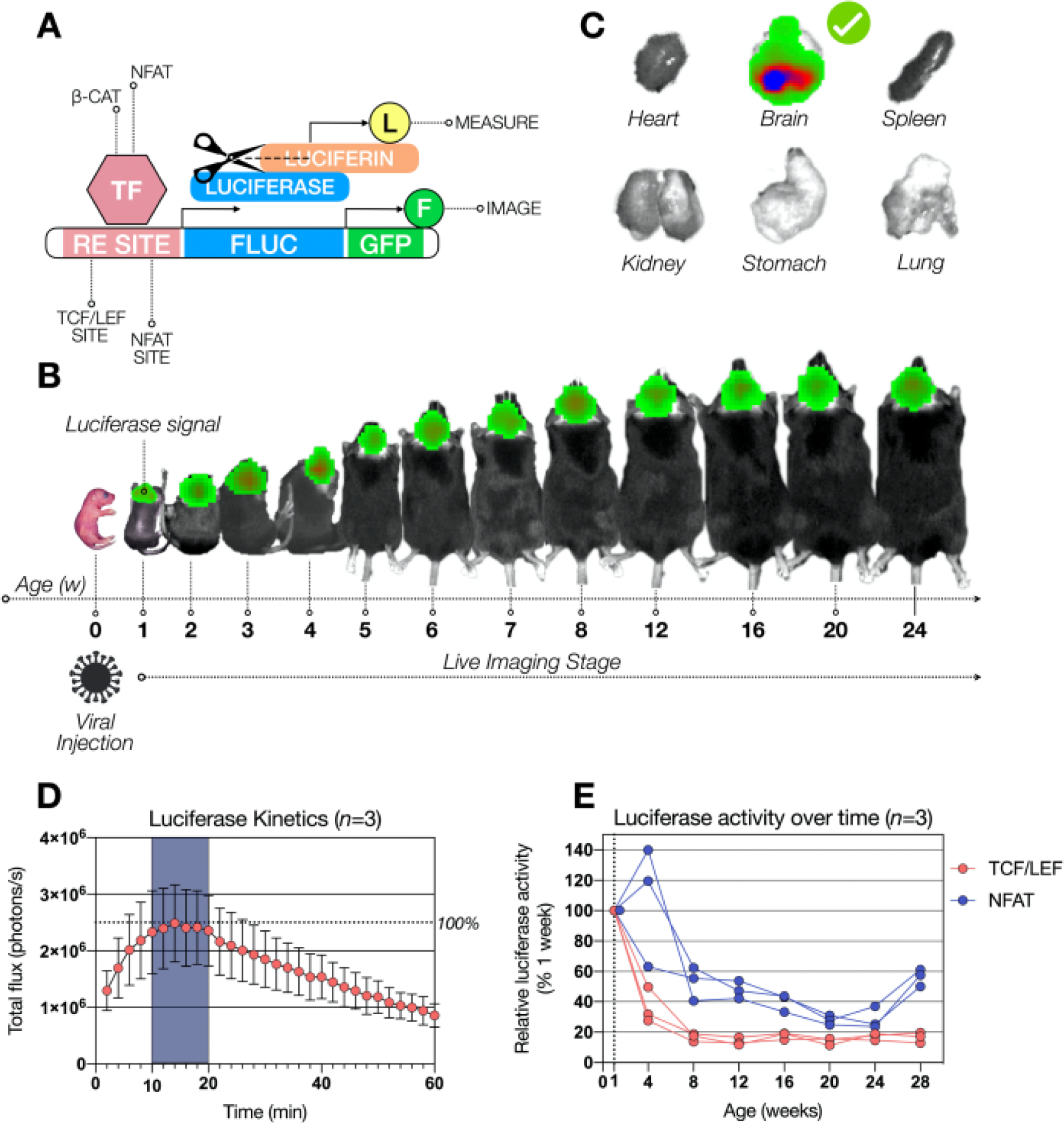
Experimental design for *in vivo* analyisis of canonical Wnt and NFATc1 signaling changes in wild type, LRRK2 KO and LRRK2 G2019S KI mice over time. Mice were injected intracranially at P0 with lentivirus containing a construct composed of a RE site for either TCF/LEF or NFATc1, a gene encoding luciferase and a second gene encoding for GFP. Expressed luciferase is able to cleave luciferin as a substrate, producing a detectable bioluminescence signal (A). This bioluminescence was measured regularly over 28 weeks in wild type (WT), LRRK2 knockout (KO) and LRRK2 G2019S knock-in (KI) mice *in vivo* (B). After 28 weeks, post-mortem analysis revealed luciferase activity prominently in the brain, excluding the other organs (C). Luciferase kinetics was determined with an ideal measurement time frame, highlighted in dark blue, n=3 (D). Relative luciferase activity is shown in WT mice over time with signal correction by the signal generated SFFV construct, used as positive control (E).

### Real time polymerase chain reaction and vector copy number analysis

To determine vector copy numbers, DNA was isolated from lentiviral injected mouse brains using the DNeasy kit from Qiagen according to the manufactures protocol with the help of a tissue grinder (Dixon Science). Relative amounts of proviral DNA have been determined by real time PCR using a StepOnePlus Real-Time PCR System (Life Technologies Limited). PCR reactions were carried out in 96 well plates following a standard thermal cycling protocol with an overall reaction volume of 20µl, containing 10µl iTaq Universal SYBR Green Supermix (Bio-Rad Laboratories Ltd.), 3µl of forward and reverse primer mix, 2µl of DNA in appropriate concentrations and 5µl of dH_2_O. Intron-spanning, target specific primers were selected with Oligo 6.71 software (MedProbe) and optimized as described previously (51). Serial dilutions of vector plasmid containing samples or wild type controls were used to generate a standard amplification curve for primer efficiency testing. All samples were tested in triplicates and PCR products were analyzed by gel electrophoresis and melting curve analysis. For vector copy number determination, the vector plasmid specific *WPRE* gene was detected and the housekeeping genes *Gapdh,* β*-actin* and *Titin* were chosen as control (49)(51). To analyze gene expression in LRRK2 knockout, LRRK2 G2019S knock-in and wild type mice, RNA was isolated from half brain, cortex, hippocampus, striatum and olfactory bulb with the RNeasy kit (Qiagen) and a tissue grinder. The isolated RNA was reverse transcribed into cDNA using the SuperScript III First-Strand Synthesis System (Life Technologies Limited). Intron-spanning, optimized primers for all tested target genes with information about sequence and product size are listed in table 1. Relative expression changes were calculated using the ΔΔCt method, correcting for primer efficiency and two housekeeper genes (*Gapdh* and *Hprt*) (52). Primer information for quantitative real time polymerase chain reaction is given in Table 1, including target gene name, accession number, primer sequence for the forward (F) and reverse (R) primer and the PCR product size to expect in genomic, copy or vector DNA. *Gapdh, Hprt,* β*-actin* and *Titin* served as housekeeper reference genes. Primers were selected with the Oligo 6.71 software, except for β-actin (51).

### Immunohistochemistry

Immunohistochemistry staining of brain slices was executed as previously described (53). Brains from PBS perfused mice were fixed in 4% paraformaldehyde for 48 hours and transferred to a 30% sucrose solution for storage at 4°C. Coronal sections of the perfused mouse brains were cryosectioned into 40μm at −20C using a Leica CM3050 cryostat (Leica Biosystems). In order to detect GFP expression via immunohistochemistry staining, slices were first incubated for 30 minutes in 1% H_2_O_2_ in Tris-buffered saline (TBS) to block endogenous peroxidase activity, followed by a three times washing step in TBS. Slices were incubated for 30 minutes in blocking solution, containing 15% normal goat serum (Sigma) in TBS with 0.3% Triton X-100 (TBS-T). Primary antibody incubation against GFP (1:10000, ab290, Abcam) happened overnight in 10% normal goat serum in TBS-T at 4°C on a rocking table. The brain slices were washed for three times in TBS and incubated in 10% normal goat serum in TBS-T with a biotinylated secondary antibody from goat against rabbit IgG (1:1000, Vector Laboratories) at room temperature for 2 hours. In order to visualize the immune staining Vectastain avidin-biotin solution (ABC, Vector Laboratories) and DAB (Sigma) were employed. Slices were mounted onto chrome gelatine coated Superfrost-plus slides (VWR), dehydrated, cleared with histoclear (National Diagnostics) for 30 minutes and coverslipped with DPX mounting medium (VWR). Non-overlapped slice images were taken under constant light intensity with a live video camera (Nikon) mounted onto a Nikon Eclipse E600 microscope. Signal intensity was finally analyzed with the help of Image-Pro Premier (Media Cybernetics) software.

### Western blot analysis

Brains from LRRK2 knockout, LRRK2 G2019S knock-in and wild type control mice were dissected into half brain, cortex, hippocampus and striatum, snap frozen in liquid nitrogen and stored at −80°C. In order to isolate total protein from these brain samples, tissue was slowly defrosted on ice, suspended in protein lysis buffer containing 5 mM MgCl_2_, 150 mM NaCl, 50 mM Tris, pH 7.5, 1% Igepal, 1x complete protease inhibitor cocktail (Roche) and 1:100 phosphatase inhibitor cocktail (Pierce), and homogenized with a tissue grinder (Dixon Science). The resulting protein lysate was centrifuged for 10 minutes at 4°C at maximum speed and the clear supernatant was transferred to a fresh collection tube. Protein concentration was determined with the bicinchoninic acid (BCA) assay (Pierce). Equal amounts of total protein were mixed with 4X NuPAGE LDS sample buffer and 10X NuPAGE reducing agent and subsequently denatured for ten minutes at 95°C with a thermo mixer and proteins were separated by size using 4-12% Bis-Tris Plus Gels (Life Technologies) in a SDS Polyacrylamide Gel Electrophoresis System (Life Technologies). Separated proteins were blotted on a polyvinylidine fluoride (PVDF) membrane with a Trans-Blot Turbo Transfer System (Bio-Rad). The protein coated membrane was blocked with 5% normal serum (Sigma) and 5% skim milk (Sigma) in Tris-buffered saline with 0.1% Tween 20 (TBS-T) for one hour, and then incubated in primary antibody solution overnight at 4°C. Primary antibodies against LRRK2 (MJFF2, Abcam), Wnt3a (Abcam), Wnt5a (Abcam), pLRP6 (S1490; Cell Signaling Technology), LRP6 (Cell Signaling Technology), pGSK-3β (pY216; BD Biosciences), GSK-3β (Cell Signaling Technology), active β-catenin (Millipore), β-catenin (New England BioLabs), TCF/LEF family antibody sampler kit (New England BioLabs), NFAT1 (New England BioLabs), BDNF (Cell Signaling Technology) and β-actin (Sigma) were diluted at desired concentration with fresh blocking buffer. On the next day, membranes were washed in TBS and incubated with HRP-conjugated secondary anti-mouse and anti-rabbit antibodies (Jackson Laboratories) in blocking buffer without normal serum for two hours on a rocking table. Membranes were washed again with TBS and signal detection was carried out with SuperSignal West Pico Chemiluminescent Substrate or SuperSignal West Femto Chemiluminescent Substrate (Pierce) and a Syngene GeneGnome Imaging system. Signal intensity was analyzed with the help of the ImageJ software.

### Primary cell culture

In this study primary cultures were generated from LRRK2 knockout, LRRK2 G2019S knock-in and wild type control mice to identify the impact of LRRK2 on Wnt and NFATc1 signaling in individual cellular systems. Primary hippocampal (54)(55) and cortical (56)(57) neurons were cultured with minor modifications according to protocols previously published. Mice were time mated to guaranty primary cells from all different genotypes at the same time. Cortical cultures were isolated from embryonic day 16 old mice and hippocampal neurons were generated from newborn (P0-1) mice. For hippocampal neuron cultures, the hippocampus form 6-8 neonatal mice was collected from both hemispheres in Hank’s buffered saline solution (HBSS, Life Technologies). Tissue was incubated in 0.25% trypsin (Life Technologies) for 10 minutes at 37°C to allow digestion. The reaction was stopped by adding trypsin inhibitor (Sigma). Four trituration steps with Neurobasal medium supplemented with B-27 (Life Technologies) led to further tissue dissection. Cells were finally plated on poly-L-lysine (Sigma) coated 24 well plates in growth medium containing Neurobasal supplemented with 1x B-27, 1% GlutaMAX, 1x N2 and 0.5% penicillin/streptomycine (all from Life Technologies). Cortical cells were isolated from 6-8 embryos per genotype in a similar way as hippocampal cultures, the final growth medium consists of Neurobasal medium supplemented with 10% GlutaMAX, 1x B-27, 1% Glucose and 0.5% penicillin/streptomycin. Cells were plated on poly-L-lysine and laminin (Sigma) coated 24 well plates. All cultures were maintained under stable conditions in a standard 37°C, 5% CO2 cell culture incubator (Binder).

### Treatment and bioluminescence imaging of cells

Primary cortical and hippocampal neurons were seeded in 24 well plates at suitable densities and transduced with lentivirus containing the following constructs: pLNT-TCF/LEF or NFAT-secNanoLuc-2A-GFP together with pLNT-SFFV-secVLuc as control at an MOI of one for two days. Transcription factor activated luciferase was expressed and secreted luciferase activity was measured in the medium 48 hours after transduction. 10µl of luciferase containing culture medium was transferred into a fresh 96 well plate and incubated with either Furimazine (Nano-Glo™ Luciferase Assay, Promega)/Coelenterazine (Prolume) in order to activate the NanoLuc or with Vargulin (Cypridina Luciferin, Prolume) in order to activate VLuc. Bioluminescence was measured with a GloMax Explorer System (Promega) using the preinstalled NanoLuc-Luciferase Activity Measurement protocol. To stimulate the cells, all cell types were treated with either 200ng Wnt3a or Wnt5a (both R and D Systems). Bioluminescence was detected 24 hours after treatment.

### Statistical analysis

Statistical significance was generally determined throughout the study by using the Prism 4.02 software (GraphPad). To analyze the *in vivo* bioluminescence imaging of mice over time, we performed a mixed-effects analysis to identify significant differences between the three LRRK2 genotypes and to test for the influence of time, involving all repeated measures over 28 weeks. The effect of sex differences was investigated by an unpaired t-test with Welch’s correction. The same test was used to identify statistical differences for genotypes in real time polymerase chain reaction experiments and western blot analysis. To analyze the *in vitro* bioluminescence imaging of cultured primary cells, we executed a 2-way ANOVA and Turkey’s multiple comparison test, determining the effect of treatment and we analyzed the effect of *LRRK2* genotype in treated and untreated primary cell cultures by using a 2-way ANOVA with Bonferroni’s multiple comparison test.

## Results

### Wnt and NFATc1 signaling are altered by LRRK2 *in vivo*

Aiming to investigate the effect of LRRK2 on Wnt and NFAT signaling, we monitored both pathways in LRRK2 knockout (KO) and LRRK2 G2019S knock-in (KI) mice in comparison to wild type (WT) mice *in vivo*. We therefore injected postnatal day 0 (P0) mice intracranially with a lentiviral construct, visualizing Wnt and NFATc1 signaling activity in a TCF/LEF and NFATc1 dependent manner. The lentiviral construct contained a response element (RE) for either TCF/LEF or NFATc1 and luciferase expression was induced through binding of the corresponding, endogenous transcription factors. Luciferin as substrate was administered to mice via intraperitoneal injection, leading to a measurable bioluminescence signal (Figure 1A). The same animals were monitored over the time frame of their first 28 weeks followed by post-mortem analyses of the organs (Figure 1B and C). *In vivo* measurements of heart, kidney, stomach, lung, spleen, and brain post-mortem verified Wnt and NFATc1 signaling activity exclusively in the brain (Figure 1C). Maximal luciferase activity was detected at 10 to 20 minutes after luciferin injection, indicating a 10-minutes window for optimal quantification (Figure 1D). Absolute quantification of Wnt and NFATc1 signaling activity over time in WT mice showed an 80% decrease of Wnt signaling activity over the first eight weeks after birth (Figure 1E). NFATc1 signaling, on the other hand increased by 40% in the first four weeks and drastically decreased to 50% after eight weeks with a further drop down to 20% after 20 weeks. The signal bounced back to approximately 50-60% at the end of the experiment.

We then investigated sex dependent canonical Wnt and NFATc1 signaling differences between LRRK2 genotypes over time (Figure 2A-G, 3A-G). Mixed-effect analysis of all mice over 28 weeks revealed a significant ∼2.9-fold (KO) and ∼2.6-fold (KI) increase in Wnt signaling activity *in vivo* (*P* < 0.0001; Figure 2A). Segregating the mice by sex exposed a 4.75-fold increase of Wnt signaling activity in male LRRK2 KO mice and a 4.66-fold increase of Wnt signaling activity in male LRRK2 G2019S KI mice with both effects being highly significant (*P* < 0.0001; Figure 2B). In female mice Wnt signaling activity was also enhanced in LRRK2 KO and LRRK2 G2019S KI mice by 1.96-fold (*P* < 0.0001) and 1.45-fold (*P* < 0.05) compared to WT mice (Figure 2C). Canonical Wnt signaling activity was more affected in male than female mice. This observation is supported by the finding that male LRRK2 WT mice show no significant difference to female LRRK2 WT mice (*P* = 0.34; Figure 2G). Time alone did not have any significant effect on signaling activity (Figure 2 D-F), showing that the above differences are reliably genotype-specific. Analysis of active β-catenin levels via Western blot at the end of the IVIS experiments in 8 randomly selected male and female KO, KI and WT mice confirmed a significant increase in canonical Wnt signalling in LRRK2 mutant KO and KI mice validating our lentiviral biosensor approach (Figure 2 H,K).

**Figure 2.**
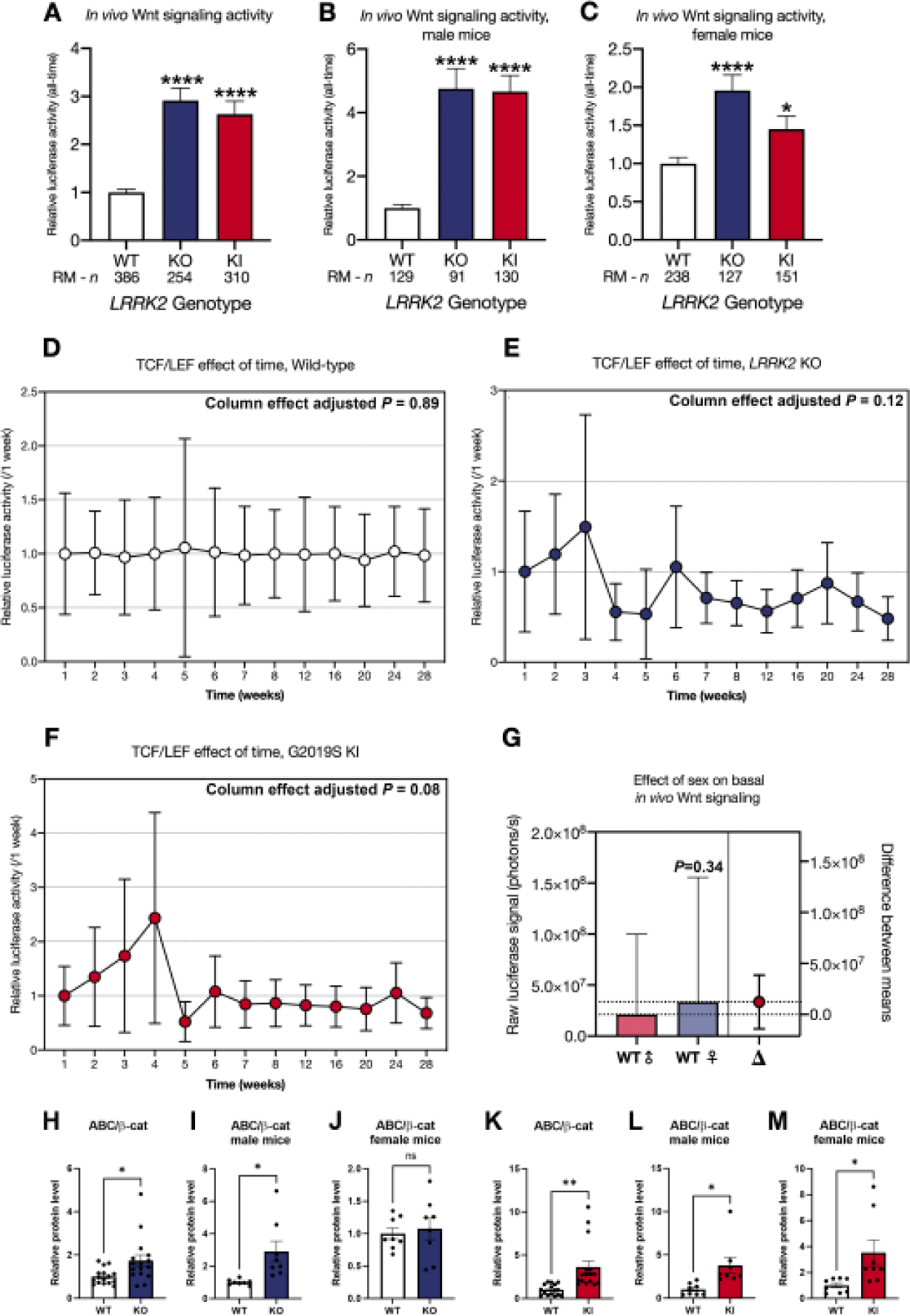
Canonical Wnt signaling changes in wild type, LRRK2 KO and LRRK2 G2019S KI mice *in vivo* over time. Mice were injected intracranially at P0 with a lentiviral biosensor for TCF/LEF to detect Wnt signaling activities. Repeated measures of the resulting bioluminescence signal were detected over a period of 28 weeks in wild type (WT), LRRK2 knockout (KO) and LRRK2 G2019S knock-in (KI) mice. Data is shown as mean relative luciferase activity ±SEM, normalized to WT set to one (**A**, **B**, and **C**). Sex separation reveals differences between *LRRK2* genotypes for only male (**B**) and only female mice (**C**). The effect of time is shown as mean relative luciferase activity ±SEM, normalized to week one set to one for each individual *LRRK2* genotype (**D**, **E**, and **F**). Sex related differences are represented as mean raw luciferase signal ±SEM in photons per seconds (**G**). Wnt signaling changes in WT, KO and KI half brain protein samples are shown as relative protein level for active β-catenin (ABC) in relation to total β-catenin (**H-M**). Data is shown as fold-change to WT and β-actin served as loading control. Statistical significance, indicated as \*\*\*\**P* < 0.0001 and \**P* < 0.05 was tested for genotype differences and effect of time by mixed-effects analysis, and the impact of sex on basal *in vivo* Wnt signaling was analyzed by unpaired t-test with Welch’s correction. Immunoblot experiments were tested via unpaired t-test with Welch’s correction with n=16, 8 males and 8 females.

Conversely, NFATc1 signaling activity was significantly reduced 0.56-fold in KI mice (*P* < 0.0001; Figure 3A). Similar effects were still observable in sex-segregated datasets (Figure 3B and 3C). NFATc1 signaling was overall not significantly affected in KO mice (*P* = 0.14; Figure 3A). Upon sex segregation, male NFATc1 signaling activity was comparable to controls (*P* = 0.16; Figure 3B), whilst female mice displayed a significant 0.69-fold decrease (*P* < 0.005; Figure 3C). Overall, no differences in Wnt and NFATc1 signaling activity were observed between male and female WT mice (*P* = 0.74; Figure 3G). Time alone did not have any significant effect on individual NFATc1 signaling measurement time points, except for KO mice, which showed a modest effect of time (Column effect adjusted *P* = 0.03; Figure 3D-F).

**Figure 3.**
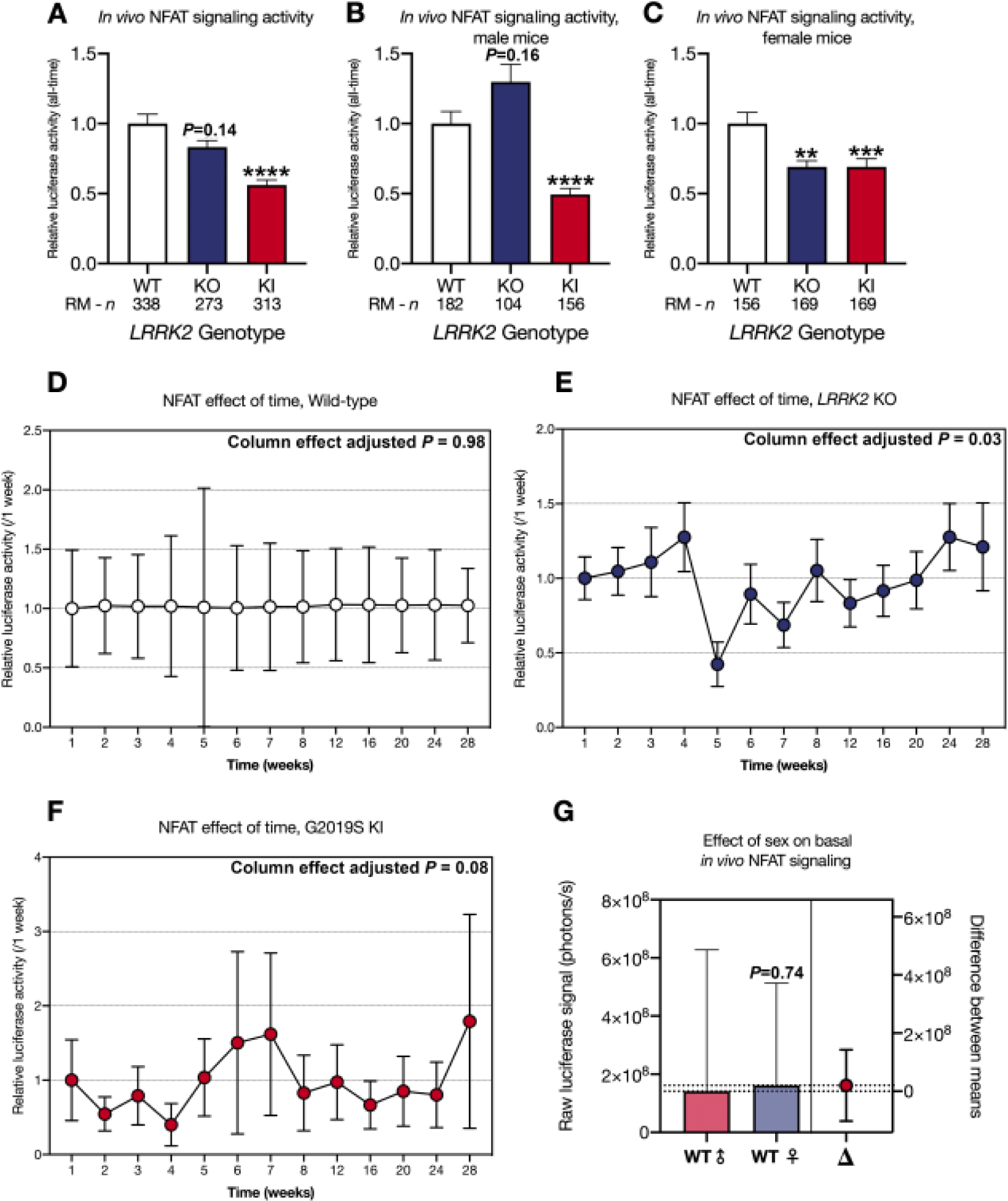
NFATc1 signaling changes in wild type, LRRK2 KO and LRRK2 G2019S KI mice *in vivo* over time. Mice were injected intracranially at P0 with a lentiviral biosensor for NFATc1 to detect NFATc1 signaling activities. Repeated measures of the resulting bioluminescence signal were detected over a period of 28 weeks in wild type (WT), LRRK2 knockout (KO) and LRRK2 G2019S knock-in (G2019S) mice. The data is shown as mean relative luciferase activity ±SEM, normalized to WT set to one (**A**, **B**, and **C**). Sex separation reveals differences between *LRRK2* genotypes for only male (**B**) and only female mice (**C**). The effect of time is shown as mean relative luciferase activity ±SEM, normalized to week one set to one for each individual *LRRK2* genotype (**D**, **E**, and **F**). Sex related differences are represented as mean raw luciferase signal ±SEM in photons per seconds (**G**). Statistical significance, indicated as \*\*\*\**P* < 0.0001, \*\*\**P* < 0.0005, and \*\**P* < 0.005 was tested for genotype differences and effect of time by mixed-effects analysis, and the impact of sex on basal *in vivo* NFATc1 signaling was analyzed by unpaired t-test with Welch’s correction.

The IVIS is limited to whole-brain resolution. Anticipating this limitation, we included a GFP marker in the lentiviral construct (Figure 1A) seeking to expand on the above observed signaling dysregulation by determining which specific brain regions were responsible for the quantified signals.

Wnt signaling activity was detectable in cells of the olfactory bulb, cortex, hippocampus, superior colliculus, striatum, pre thalamus, and thalamus (Figure 4). We found NFATc1 signaling activity in the same brain regions, excluding the striatum (Figure 5). Positive staining for Wnt and NFATc1 signaling was further evident in the inferior colliculus, dorsal lateral geniculate nucleus (dLGN), hypothalamus, and lateral ventricle (Figure S1 and S2). We detected no GFP positive cells in the brain of non-injected negative control mice (Figure 4, 5, S1 and S2).

**Figure 4.**
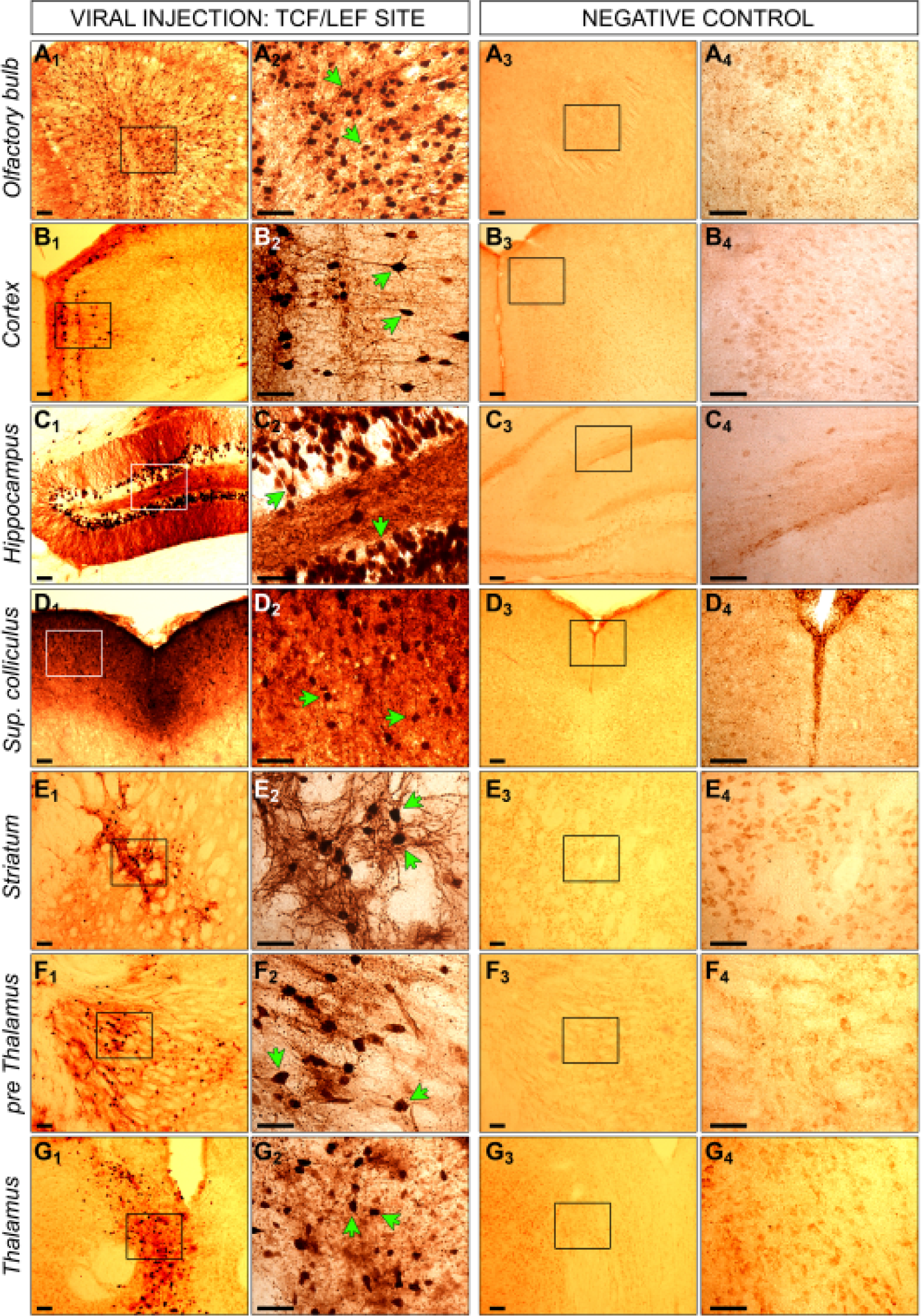
GFP staining of injected and non-injected mice with the TCF/LEF biosensor. Mice were injected intracranially at P0 with a lentiviral construct, containing a RE site for TCF/LEF and a second gene for GFP. Six months later, brains were collected from injected (**1** and **2**) and non-injected control mice (**3** and **4**). Brains were fixed in PFA, coronally cryosectioned into 40µm thick slices, which were stained via immunohistochemistry for GFP. Positive cells for GFP expression are visible in dark brown indicated by the green arrows. Positive cells were detectable in olfactory bulb (**A**), cortex (**B**), hippocampus (**C**), superior colliculus (**D**), striatum (**E**) and thalamus (**F** and **G**). No positive cells were measurable in corresponding regions of non-injected control brains. Images in **2** and **4** represent the zoom of indicated squares in **1** and **3**. Scale bar for all images is 50µm.

**Figure 5.**
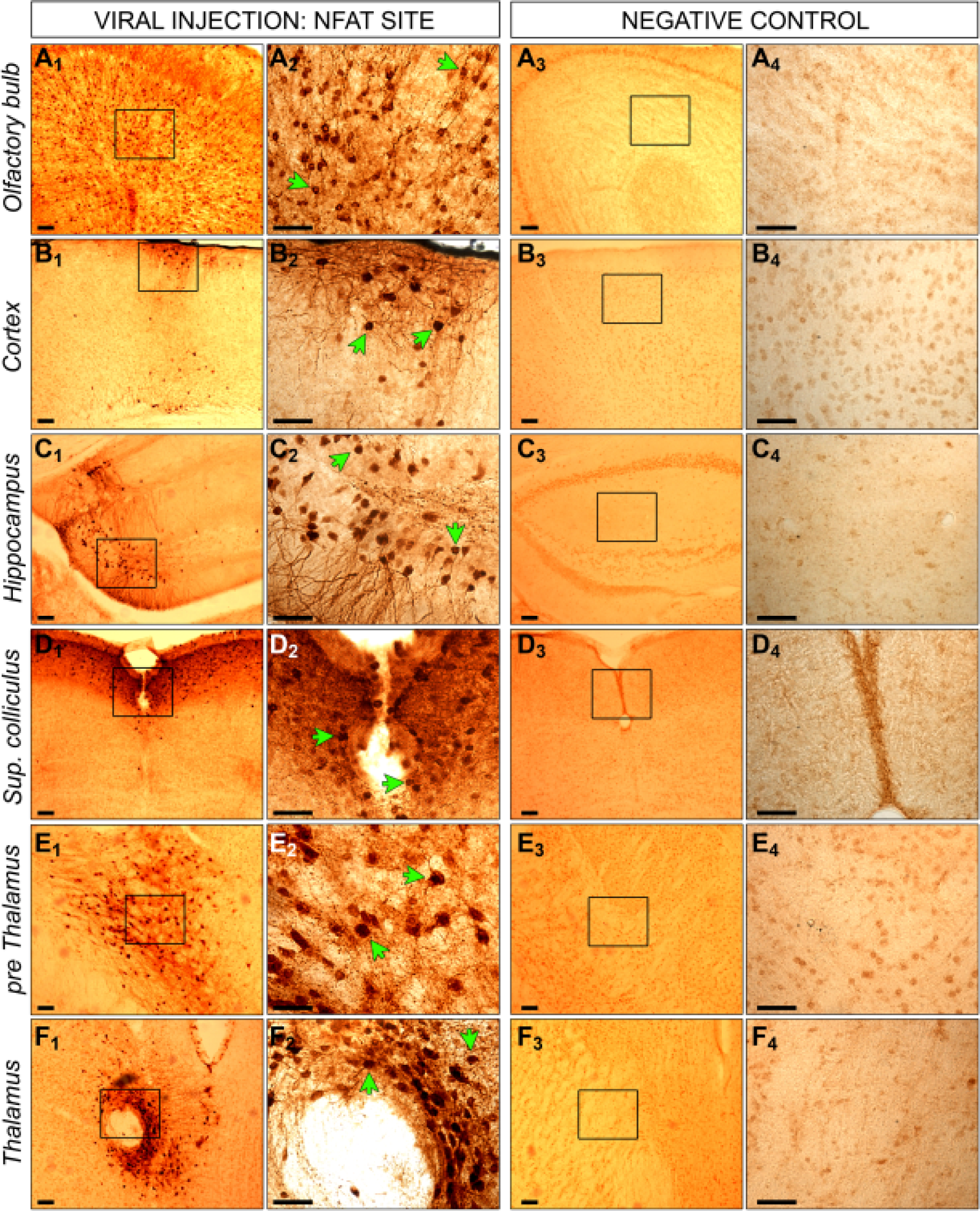
GFP staining of injected and non-injected mice with the NFATc1 biosensor. Mice were injected intracranially at P0 with a lentiviral construct, containing a RE site for NFATc1 and a second gene for GFP. Six months later, brains were collected from injected (**1** and **2**) and non-injected control mice (**3** and **4**). Brains were fixed in PFA, coronally cryosectioned into 40µm thick slices, which were stained via immunohistochemistry for GFP. Positive cells for GFP expression are visible in dark brown indicated by the green arrows. Positive cells were detectable in olfactory bulb (**A**), cortex (**B**), hippocampus (**C**), superior colliculus (**D**) and thalamus (**E** and **F**). No positive cells were measurable in corresponding regions of non-injected control brains. Images in **2** and **4** represent the zoom of indicated squares in **1** and **3**. Scale bar for all images is 50µm.

Based on these observations, we elected to focus our subsequent investigations on the following GFP-positive regions: cortex, hippocampus and striatum, seeking to further characterize any potential LRRK2-mediated WNT and NFAT dysregulation at the gene expression and protein levels.

### Wnt and NFAT signaling changes by LRRK2 are tissue specific

To explore the physiological and pathogenic function of LRRK2 on a molecular level, we combined qPCR and immunoblotting to assess gene expression and protein level of selected relevant Wnt and NFAT signaling components in LRRK2 KO and G2019S KI in comparison to WT mice in cortex, hippocampus, striatum and half brain samples. Our choice of explored signaling components was led by tested qPCR primer efficacy, availability of reliable antibodies and our primary focus on canonical Wnt signaling. Canonical Wnt signaling mediators investigated were the LRP6 co-receptor, Wnt3a and Wnt7a receptor ligands, the cytosolic β-catenin destruction complex proteins Axin2 and GSK-3β, the transcription factors β-catenin, TCF and LEF, as well as the Wnt target gene BDNF. Wnt5a was investigated as a non-canonical Wnt signaling ligand opposing canonical Wnt signaling, and the transcription factor NFATc1 as a correlate for NFAT signaling.

We assumed that changes in half brain might reflect best the signaling changes observed *in vivo*. Confirming our *in vivo* observations of an increase in canonical Wnt signaling by mutant LRRK2 on a molecular level we showed that active β-catenin/total β-catenin was increased significantly in KO and G2019S KI mice in our IVIS mouse cohort at 28 weeks of age. This effect also reached significance in male KO, and male and female G2019S KI mice, whereas the effect did not reach significance for female KO mice (Figure 2H-M). In half brain of LRRK2 KO mice we also detected significant downregulation of *Wnt5a,* β*-catenin, Axin2,* and *Gsk-3*β (Figure 6A_1_) mRNA levels while the protein level of BDNF was significantly increased and TCF1 trended towards an increase (*p* = 0.057; Figure 6B_1_). These changes are overall suggestive of canonical Wnt signaling upregulation. In addition, and also in line with the canonical Wnt signaling upregulation observed *in vivo,* LRRK2 G2019S KI half brains presented significantly increased relative *Wnt7a* mRNA levels and a trend towards decreased *Wnt5a* gene expression (*p* = 0.06; Figure 6A_2_). These signaling changes are further underpinned by a significant increase of TCF1 protein levels in G2019S KI brains (Figure 6B_2_), whilst Wnt5a levels were significantly increased, we observed a stark decrease in BDNF and pLRP6 (S1490) (Figure 6B_2_).

**Figure 6.**
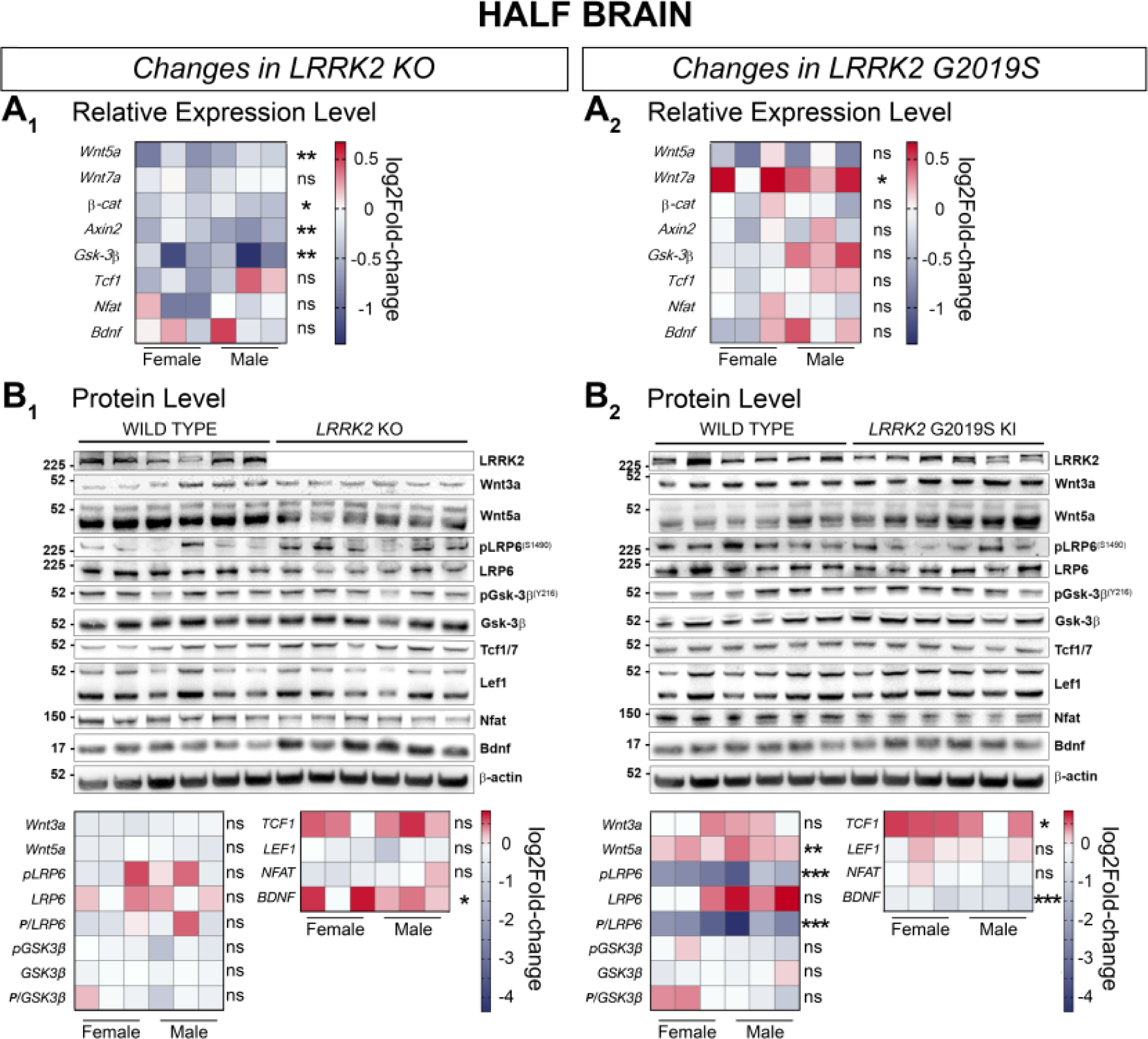
Signaling pathway component changes in the half brain of LRRK2 KO and LRRK2 G2019S KI mice compared to WT mice. Wnt and NFAT signaling component changes in half brain samples between LRRK2 KO and WT (**A1, B1**), and LRRK2 G2019S KI and WT (**A2, B2**) are shown on a transcriptional (**A**) and protein level (**B**). Relative expression of relevant gene candidates was detected via quantitative real time PCR. Data is shown as log_2_fold-change to WT expression. *Gapdh* and *Hprt* served as housekeeping genes (**A_1_, A_2_**). Statistical significance shown as \*\**P* < 0.005 and \**P* < 0.05 was tested via unpaired t-test with Welch’s correction with n=6, 3 males and 3 females (**B_1_**, **B_2_**). Representative plots are displayed, and data is shown as log_2_fold-change to WT protein levels. β-actin served as loading control. Statistical significance indicated as \*\*\**P* < 0.0005, \*\**P* < 0.005, and \**P* < 0.05 was tested via unpaired t-test with Welch’s correction with n=6, 3 males and 3 females.

In cortical samples of LRRK2 KO mice we found significantly decreased protein levels of Wnt3a and pLRP6 (S1490), both standalone and as a fraction of total LRP6 (Figure 7B_2_). BDNF protein levels showed a trend towards upregulation (*P* = 0.056), while expression levels for *Bdnf* were significantly downregulated in the cortex of LRRK2 KO mice (Figure 7A_1_ and 7B_2_).

**Figure 7.**
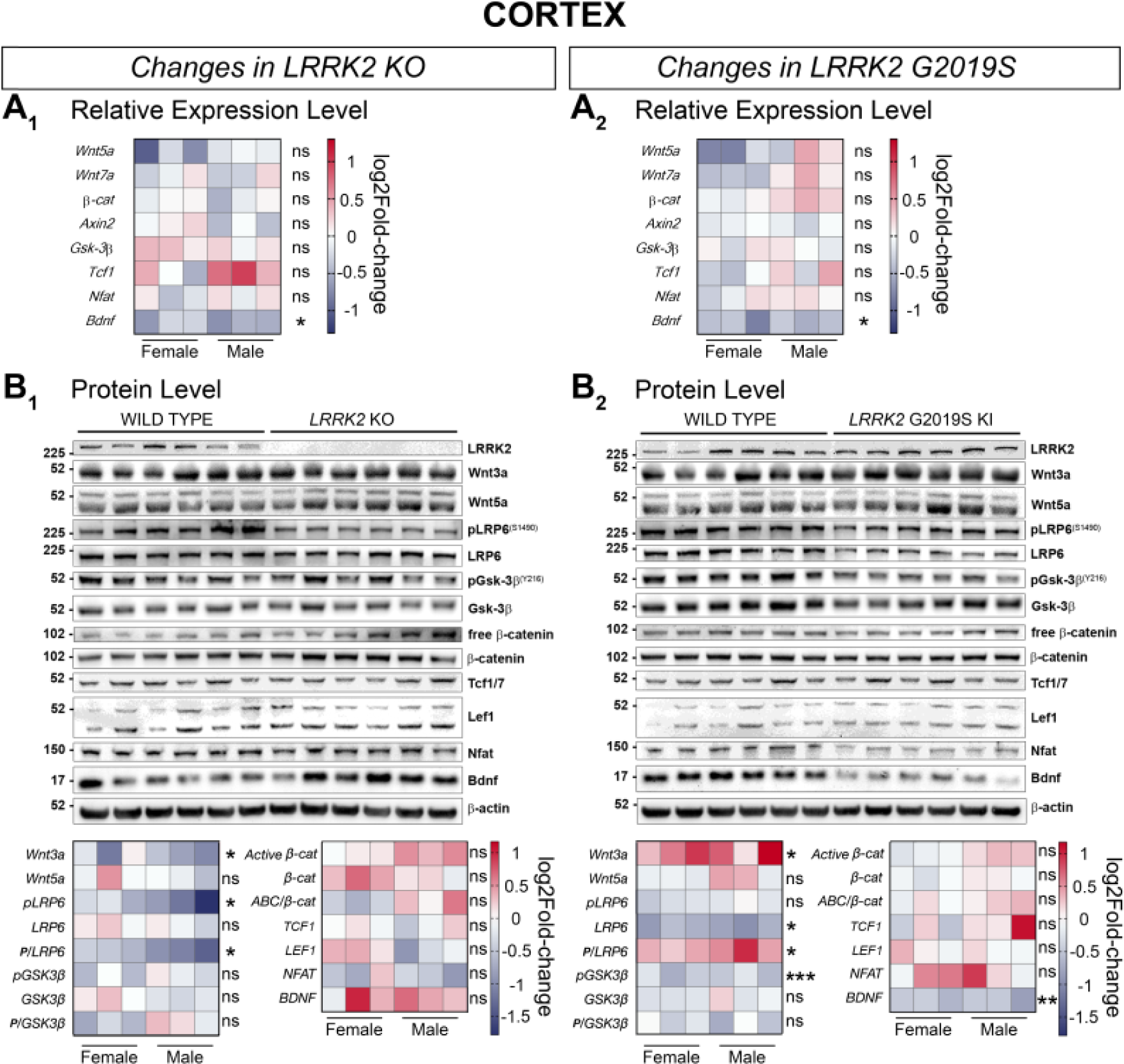
Signaling pathway component changes in the cortex of LRRK2 KO and LRRK2 G2019S KI mice compared to WT mice. Wnt and NFAT signaling component changes in in cortical samples between LRRK2 KO and WT (**A1, B1**), and LRRK2 G2019S KI and WT (**A2, B2**) are shown on a transcriptional (**A**) and protein level (**B**). Relative expression of relevant gene candidates was detected via quantitative real time PCR. Data is shown as log_2_fold-change to WT expression. *Gapdh* and *Hprt* served as housekeeping genes (**A_1_, A_2_**). Statistical significance shown as \**P* < 0.05 was tested via unpaired t-test with Welch’s correction with n=6, 3 males and 3 females (**B_1_**, **B_2_**). Representative plots are displayed, and data is shown as log_2_fold-change to WT protein levels. β-actin served as loading control. Statistical significance indicated as \*\*\**P* < 0.0005, \*\**P* < 0.005, and \**P* < 0.05 was tested via unpaired t-test with Welch’s correction with n=6, 3 males and 3 females.

In LRRK2 G2019S KI cortices, *Bdnf* gene expression was significantly decreased, whilst the expression level of other investigated gene candidates remained unchanged (Figure 7A_2_). At the protein level, significant upregulation was detected for Wnt3a and pLRP6 (S1490) in relation to total LRP6 in the cortex of KI mice (Figure 7B_2_). Total LRP6, phosphorylated GSK-3β (Y216) and BDNF protein levels were on the other hand significantly reduced in LRRK2 G2019S KI cortical samples (Figure 7B_2_).

Gene expression of relevant Wnt and NFAT signaling candidates remained largely unchanged in the hippocampus of LRRK2 KO and LRRK2 G2019S KI mice (Figure 8A_1_ and 8A_2_). Western blot analysis displayed a significantly reduced level for active β-catenin/total β-catenin and TCF1 in KO hippocampi, clearly speaking for a decrease in canonical Wnt signaling activity (Figure 8B_1_). As in the LRRK2 KO hippocampus, total β-catenin protein levels were significantly upregulated whereas TCF1 protein levels were significantly downregulated in the hippocampus of LRRK2 G2019S KI mice, (Figure 8B_2_). Despite these similarities in the expression pattern in LRRK2 KO and KI hippocampi, the ratio of β-catenin/total β-catenin did not decrease significantly in the G2019S KI mice. Nevertheless, the majority of proteins remained unchanged in the hippocampus of KO and KI mice (Figure 8B_1_ and 8B_2_).

**Figure 8.**
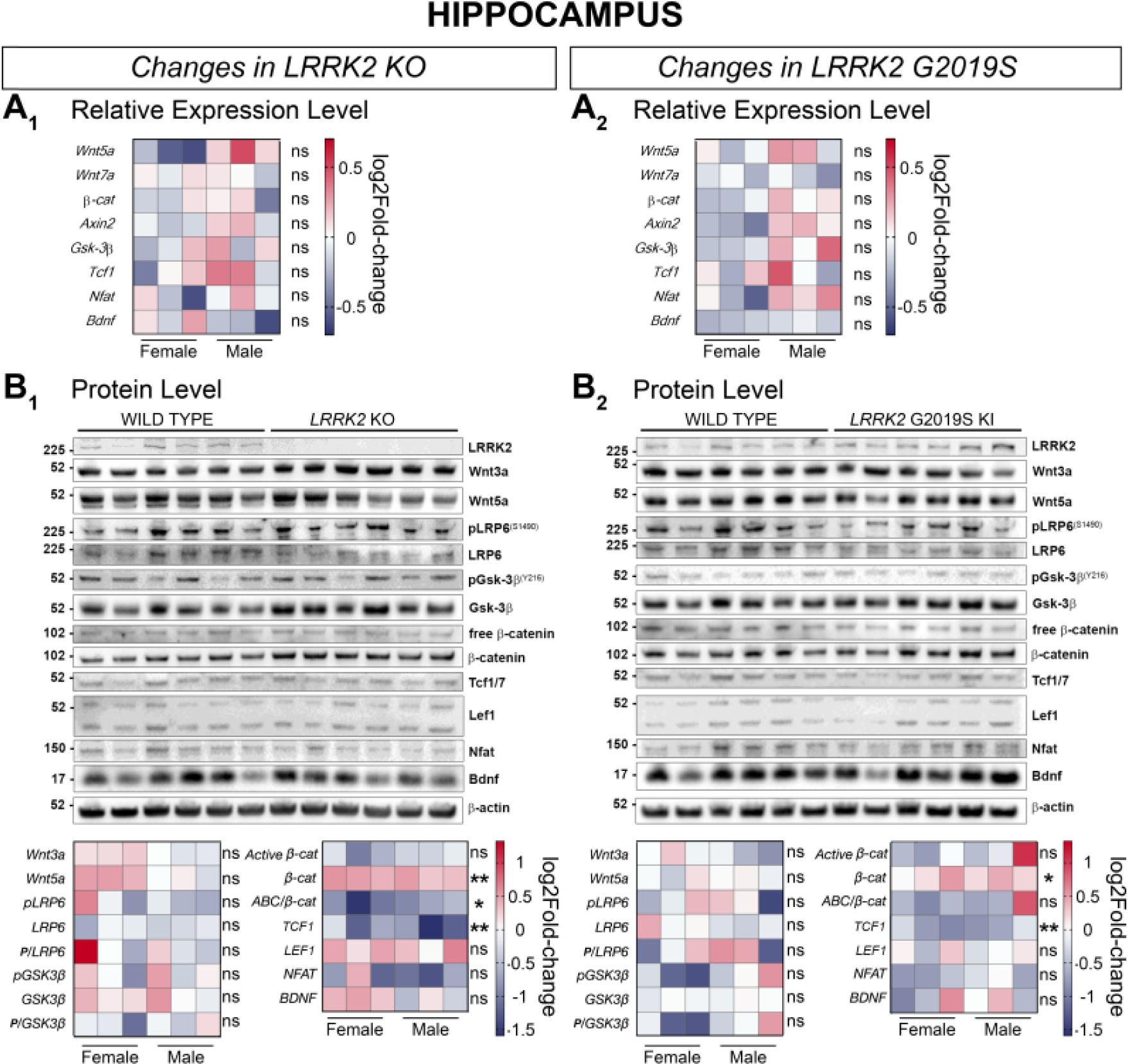
Signaling pathway component changes in the hippocampus of LRRK2 KO and LRRK2 G2019S KI mice compared to WT mice. Wnt and NFAT signaling component changes in hippocampal samples between LRRK2 KO and WT (**A1, B1**), and LRRK2 G2019S KI and WT (**A2, B2**) are shown on a transcriptional (**A**) and protein level (**B**). Relative expression of relevant gene candidates was detected via quantitative real time PCR. Data is shown as log_2_fold-change to WT expression. *Gapdh* and *Hprt* served as housekeeping genes (**A_1_, A_2_**). Statistical significance was tested via unpaired t-test with Welch’s correction with n=6, 3 males and 3 females (**B_1_**, **B_2_**). Representative plots are displayed, and data is shown as log_2_fold-change to WT protein levels. β-actin served as loading control. Statistical significance indicated as \*\**P* < 0.005, and \**P* < 0.05 was tested via unpaired t-test with Welch’s correction with n=6, 3 males and 3 females.

Most genes in male LRRK2 KO and KI striatal samples showed a trend towards upregulation in their relative expression compared to WT samples, while female mRNA expression levels remained overall unchanged (Figure 9A_1_ and 9A_2_). However, we were able to detect significant augmented gene expression for both sexes in *β-catenin, Axin2,* and *Nfatc1* in the striatum of KO mice (Figure 9A_1_). Proteins showed on the other hand a change towards downregulation in KO and KI striatal samples (Figure 9B_1_ and 9B_2_). Significant decrease was detected for Wnt5a and LEF1 in KO striatum tissue and for LRP6, phosphorylated GSK-3β (Y216) in relation to total GSK-3β, β-catenin, LEF1 and NFAT in KI samples (Figure 9B_1_ and 9B_2_). The protein level of GSK-3β was significantly increased in LRRK2 G2019S KI striatal samples (Figure 9B_2_). Active β-catenin (*P* = 0.052) showed however a tendency towards downregulation in the striatum of G2019S KI mice (Figure 9B_2_).

**Figure 9.**
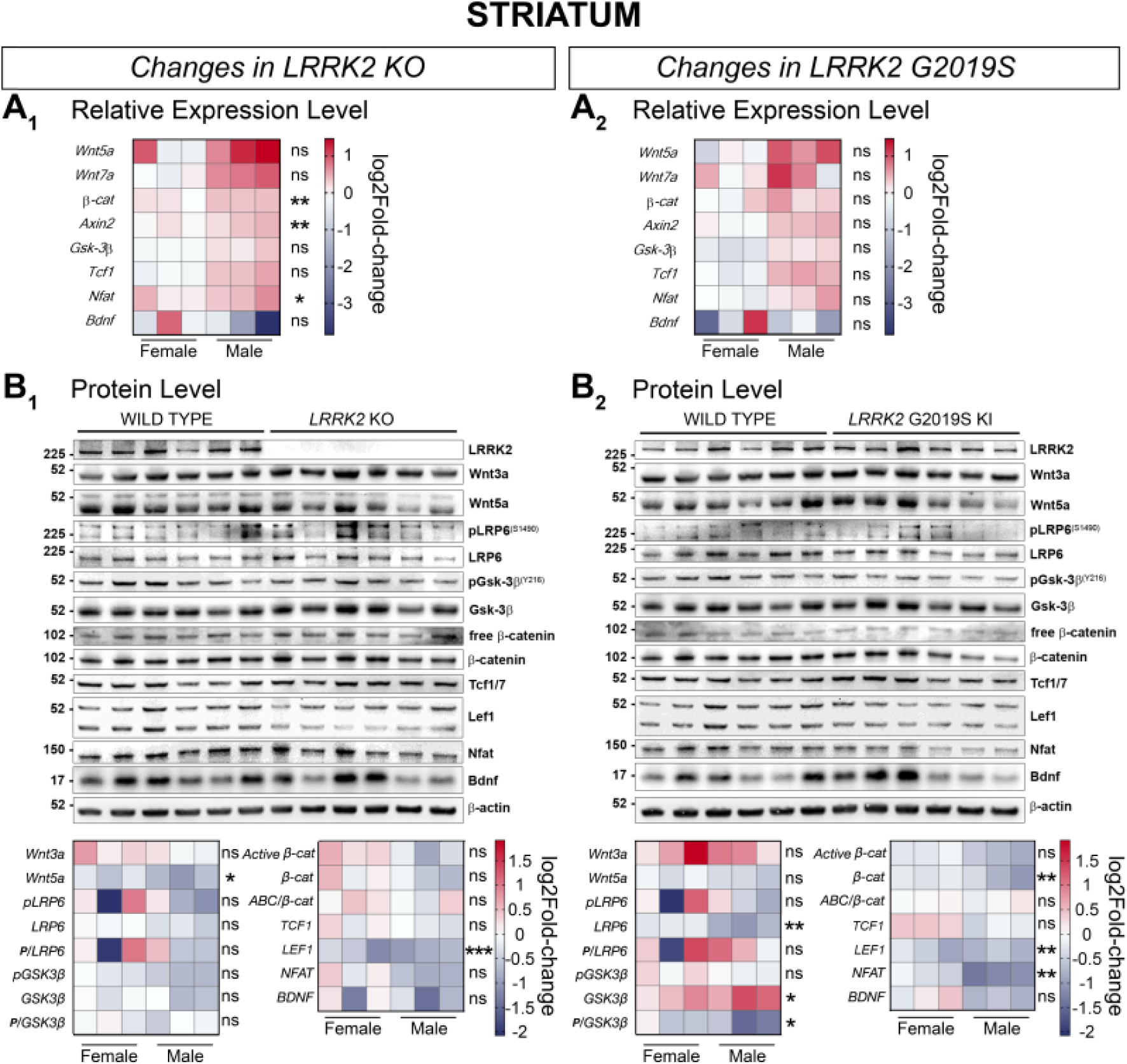
Signaling pathway component changes in the striatum of LRRK2 KO and LRRK2 G2019S KI mice compared to WT mice. Wnt and NFAT signaling component changes in striatal samples between LRRK2 KO and WT (**A1, B1**), and LRRK2 G2019S KI and WT (**A2, B2**) are shown on a transcriptional (**A**) and protein level (**B**). Relative expression of relevant gene candidates was detected via quantitative real time PCR. Data is shown as log_2_fold-change to WT expression. *Gapdh* and *Hprt* served as housekeeping genes (**A_1_, A_2_**). Statistical significance shown as \*\**P* < 0.005 and \**P* < 0.05 was tested via unpaired t-test with Welch’s correction with n=6, 3 males and 3 females (**B_1_**, **B_2_**). Representative plots are displayed, and data is shown as log_2_fold-change to WT protein levels. β-actin served as loading control. Statistical significance indicated as \*\*\**P* < 0.0005, \*\**P* < 0.005, and \**P* < 0.05 was tested via unpaired t-test with Welch’s correction with n=6, 3 males and 3 females.

### Wnt and NFAT signaling changes by LRRK2 are cell type specific

To identify cell type specific effects of LRRK2, we cultured primary cortical and hippocampal neurons from WT, LRRK2 KO and LRRK2 G2019S KI mice and adapted our *in vivo* biosensors for cell culture usage. Primary cells were transduced with Wnt and NFATc1 signaling biosensors and stimulated with the canonical Wnt ligand Wnt3a and the opposing non-canonical Wnt5a.

Wnt and NFATc1 signaling activity was increased in cortical neurons from WT and LRRK2 KO mice upon Wnt3a stimulation (Figure 10A_1_). Wnt5a did not affect Wnt and NFATc1 signaling activity in cortical KO cultures (Figure 10A_1_). However, a significant decrease in Wnt signaling activity was detected in cortical KO compared to WT cells under Wnt3a-stimulated conditions (Figure 10A_1_, left). No genotypic differences in NFATc1 signaling were observed in these cultures (Figure 10A_1_, right).

**Figure 10.**
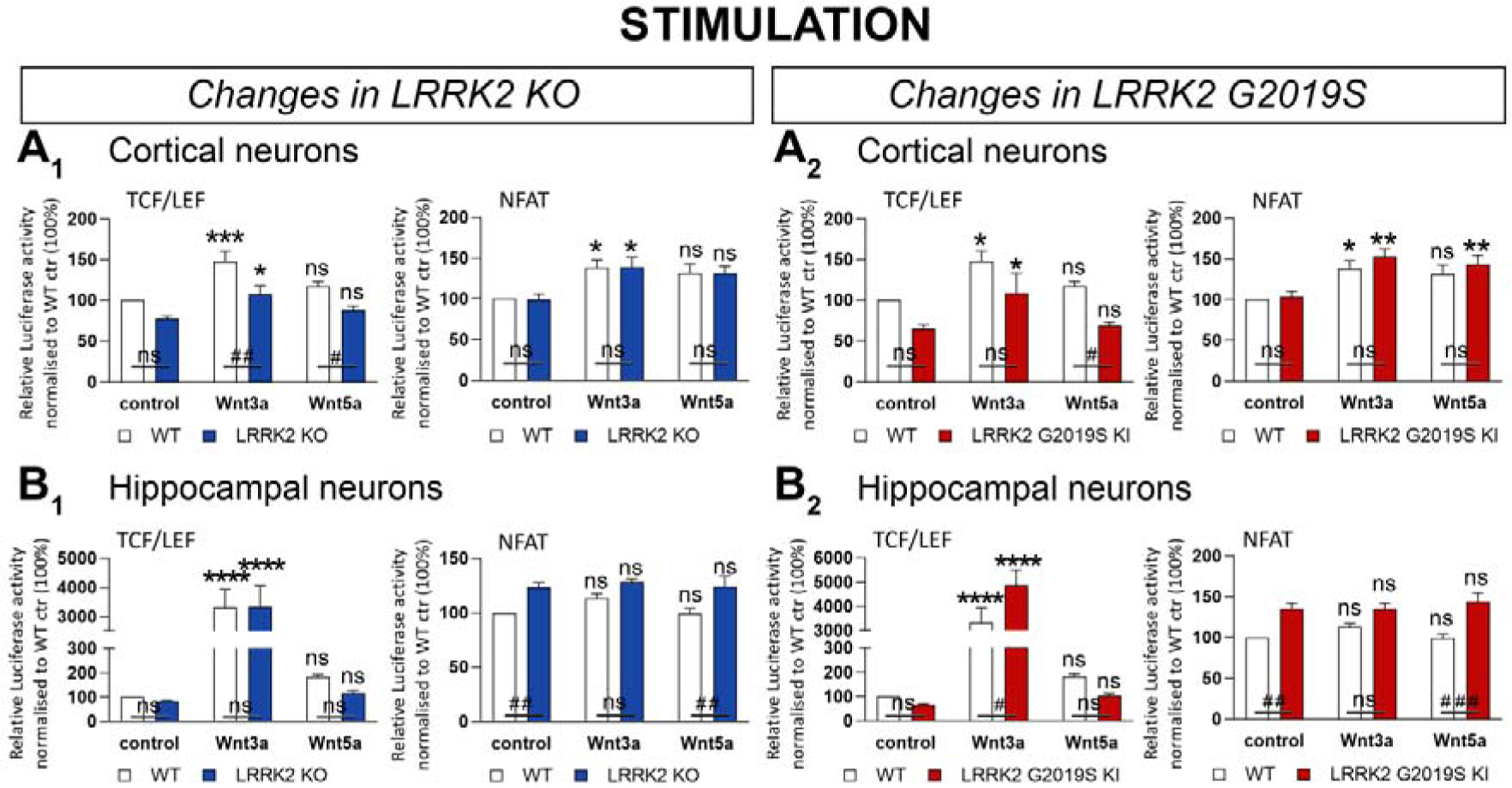
Stimulation of primary neuronal cultures from LRRK2 KO, LRRK2 G2019S KI and WT mice. Wnt (TCF/LEF) and NFATc1 signaling changes are presented under treated and untreated conditions in primary cultured cortical (**A**) and hippocampal cells (**B**) from LRRK2 KO (1), LRRK2 G2019S KI (2), and WT mice. Cells were transduced with a lentiviral biosensor displaying Wnt or NFATc1 signaling activity. Data is shown as mean relative luciferase activity ±SEM, normalized to untreated WT set to 100%. Bioluminescence was measured 24h after treatment. Statistical significance shown as ###*P* < 0.0005, ##*P* < 0.005, and #*P* < 0.05 was tested for the effect of *LRRK2* genotype via 2-way ANOVA and Bonferroni’s multiple comparison test and the effect of treatment shown as \*\*\*\**P* < 0.0001, \*\*\**P* < 0.0005, \*\**P* < 0.005, and \**P* < 0.05 was tested via 2-way ANOVA and Turkey’s multiple comparison test with n=5 independent cultures.

Wnt and NFATc1 signaling were significantly upregulated upon Wnt3a stimulation in cortical WT and KI cells (Figure 10A_2_). Wnt5a significantly enhanced NFATc1 signaling activity exclusively in cortical KI neurons (Figure 10A_2_). Wnt5a stimulated Wnt signaling activity was downregulated in cortical KI neurons compared to WT cells (Figure A_2_, left).

Wnt signaling activity was significantly enhanced by at least 30-fold by Wnt3a stimulation in hippocampal cultures across all genotypes (Figure 10B_1_ and 10B_2_, left). NFATc1 signaling remained unchanged in hippocampal cultures upon Wnt3a/5a stimulation (Figure 10B_1_ and 10B_2_, right). Neither Wnt3a nor Wnt5a activated NFATc1 signaling in hippocampal neurons, with comparable levels between basal and stimulated conditions. We did however observe a significant basal NFATc1 elevation in KO and KI cells. In addition, hippocampal KI neurons displayed significantly enhanced Wnt activation with Wnt3a but not Wnt5a stimulation in comparison to wild neurons (Figure 10B_2_).

## Discussion

In this study, we developed a lentiviral-based biosensor assay for luciferase-mediated *in vivo* monitoring of signaling pathways over time. We then employed this method to investigate the regulatory role of LRRK2 on Wnt and NFAT signaling. Although the lentiviral biosensor system provided a reliable method for comparing cell signalling activity between mutant and WT mice, we observed eGFP expression differences between brain areas such as the cortex and olfactory bulb that are not easily explained by differential cell signalling activity alone (Figures 4A-B, 5A-B). Part of this observation is reflected by the distance between brain areas and the virus ICV injection site at the lateral ventricle. In addition, the preferential distribution of lentiviruses via neural pathways such as the rostral migratory stream (RMS), physiologically used as a route for cells such as neural progenitor cells, allows enhanced viral delivery from the lateral ventricles to olfactory bulbs (58)(59)(60). However, the reliability of the biosensor signal as a readout for cell signalling activity in whole brain was validated via examining active β-catenin expression levels in half brain tissue (Figure 2H-M).

LRRK2 is a complex protein with many interaction partners. Besides its enzymatic activity, LRRK2 was previously shown to maintain homeostasis of signaling pathways by functioning as scaffolding protein (17), supporting the enzymatic activity and function of other proteins and protein complexes. Understanding the function of LRRK2 and the physiological role of pathogenic LRRK2 precisely, gained particular interest, since its linkage to PD in 2004 (8)(9). In hereditary PD, mutations in the *LRRK2* gene lead to similar symptoms as idiopathic cases. Furthering our understanding of how LRRK2 influences signaling pathways over time, such as Wnt and NFAT signaling is necessary to identify potential novel therapeutic targets. To this end, in this study we monitored Wnt and NFAT signaling activity in WT, *LRRK2* KO, and *LRRK2* G2019S KI mice right after birth *in vivo*, over a 28-week time span (Figure 1-3).

G2019S is the most prevalent PD-related *LRRK2* mutation in the general population. Canonical Wnt signaling activity was clearly upregulated in *LRRK2* KO and KI mice (Figure 2A-C), while NFAT signaling was downregulated in *LRRK2* KI mice and female *LRRK2* KO mice (Figure 3A-C). Whereas, this is our first study into NFAT signaling in LRRK2 models and canonical Wnt signaling in G2019S mice, we have previously observed a dose dependent increase in active β-catenin levels in in male and female brain tissue in homozygous and heterozygous *LRRK2* KO mice (61). An increase in canonical Wnt signaling activity has also previously been shown, in *LRRK2* KO MEF cells and upon LRRK2 knockdown in immortalized cell lines, in line with observations from other investigators showing that LRRK2 inhibits canonical Wnt signaling (6)(18)(61). Unexpectedly, *LRRK2* KO and KI resulted in the same directional change in both signaling pathways under basal conditions *in vivo*. This was in contrast to previous observations in overexpression immortalized cell systems suggestive of canonical Wnt signaling activity regulation in opposite directions (18)(62). However, we also reported previously that loss of LRRK2 increased Wnt signaling activity, while LRRK2 kinase inhibition decreased Wnt signaling activity (61). Although the underlying mechanism remains unclear, loss of LRRK2 and increased LRRK2 kinase activity might both upregulate Wnt signaling activity. Further supporting this idea, is the observed similarity in phenotype throughout our investigations in different brain regions and cell models. It is also important to note that similar phenotypes for mutant *LRRK2* and loss-of-function models have been observed previously in studies investigating the nuclear morphology of striatal projection neurons and demonstrating axon growth impairment under both conditions (62)(63). This suggests that loss of function and gain of kinase activity in LRRK2 might have a similar effect on certain cellular pathways. It is also interesting to note that a recent study reported motor impairment and loss of dopaminergic neurons in the substantia nigra pars compacta of double LRRK1/LRRK2 KO mice suggesting the relevance of LRRK2 mutant and KO to parkinsonian phenotypes (64).

Segregating the mice by sex, we observed that *in vivo* signaling changes were not homogenous between male and female animals as already observed in our previous publication, suggesting more increase in canonical Wnt signaling activity in male than female *LRRK2* KO mice, albeit in a very small number of animals (61). Wnt upregulation was more pronounced in male than female mice for both mutant genotypes (Figure 2B-C). Whilst NFAT signaling was consistently downregulated in female KO and KI mice (Figure 3C), *LRRK2* KO had no effect in male mice (Figure 3B). This result perhaps contributed to the loss of significance observed when analyzing all *LRRK2* KO animals together, regardless of sex, for NFAT signaling changes. Several sex differences are well known in PD patients. For instance, men are twice as likely as women to develop the condition (66), but women suffer from faster disease progression and increased mortality (67). Our data suggests that Wnt and NFAT activity might be differentially affected by LRRK2 modulation in males and females. However, further investigation is needed to determine the impact of this observation, if any, on the risk of PD development and its severity.

We used a two-fold approach to validate the reliability of our novel method. First, we employed immunohistochemistry to confirm effective delivery of the vector biosensor construct to individual and specific murine brain regions (Figure 4-5). GFP staining confirmed that the Wnt biosensor had been transduced effectively across all key brain areas. This result is particularly important, as it allowed us to conclude that the observed luminescence did indeed reflect *in vivo* cerebral Wnt signaling activity. Secondly, due to the novelty of this method, we deemed it essential to determine whether our *in vivo* signaling findings could be replicated in more established model systems, such as immunoblotting of active β-catenin species. For both genotypes, we observed significant Wnt upregulation (Figure 2H-M), in line with the IVIS data.

We then sought to determine how LRRK2 may cause the *in vivo* observed changes on a molecular level, by analyzing gene expression and protein level changes of specific Wnt and NFAT signaling candidates in WT, *LRRK2* KO and *LRRK2* G2019S KI mice. We assessed a high number of molecular components of known importance to Wnt and NFAT signaling (Figure 6-9). We first investigated molecular changes in half-brains, to better represent system-level signaling activity patterns. Overall, while both KO and G2019S mice showed an overall increase of canonical Wnt signaling *in vivo* and via immunoblotting of active β-catenin in half brains, specific Wnt component up-/downregulation differed markedly. This suggests different mechanisms leading to signaling activity changes in LRRK2 KO and G2019S KI mice confirming the idea that the LRRK2 G2019S mutant is not a ‘*simple*’ loss of function mutation.

In *LRRK2* KO half brain samples, the relative expression of *Wnt5a*, *Axin2*, and *Gsk-3*β was reduced. The latter two are known LRRK2 interactors (18)(68)(69), suggesting a potential functional link relating back to the observed *in vivo* phenotype. *Axin2* and *Gsk-3*β are particularly important in this respect. Given their known Wnt-suppressive functions, reduced expression of these components in *LRRK2* KO mice might account for the observed enhancement of Wnt signaling activity. Wnt5a is a non-canonical Wnt agonist, which may suppress β*-*catenin signaling (30)(31). This notion may account for the observed reduction of *Wnt5a* expression. Despite these considerations, the above mRNA findings were not reflected at the protein level. The only molecular component whose protein level was significantly affected was brain-derived neurotrophic factor (BDNF). This mediator is important in neuronal development and survival (70)(71). Previous studies suggest BDNF expression may be regulated by canonical Wnt signaling in a neuronal activity-dependent manner (72)(73). BDNF may also stimulate neuronal growth via canonical Wnt signaling through GSK-3β modulation (74). Interestingly, circulating BDNF levels are enhanced in idiopathic PD patients (75). Such patients reportedly feature longer disease duration, more severe motor deficiencies and mild cognitive loss. On the other hand, *decreased* levels of BDNF are also associated with PD as well as Alzheimer’s diseases (75)(76). We found a significant *downregulation* of BDNF protein levels in half brain samples of the pathogenic *LRRK2* G2019S KI model. In the same model, the protein level of the Wnt effector TCF1 was significantly increased, as was as the relative gene expression of *Wnt7a,* a known canonical Wnt ligand (29). These two findings overall support the *in vivo* finding of enhanced Wnt signaling activity in KI mice. However, we also observed a protein-level increase in Wnt5a and a decrease in pLRP6 (S1490) in KI half brains. LRP6 is a canonical Wnt co-receptor, important in its activation (77)(78), whilst Wnt5a may suppress the pathway (79), as mentioned. These results are thus somewhat counterintuitive, as they would more likely reflect Wnt signaling suppression but might also be the result of a negative feedback regulation.

Due to their relative importance in PD (80)(81)(82), high expression of *LRRK2* (27) as well as the reliable vector delivery observed via immunohistochemistry, we selected the striatum, cerebral cortex and hippocampus for further study via the same dual gene expression/protein level approach. To our surprise, when narrowing the scope to more specific brain areas, the molecular patterns observed were markedly different. Despite complex changes in signaling components the only brain region investigated showing a significant change in active β-catenin/total β-catenin is the LRRK2 KO hippocampus. In striatum and cortex, the signaling mediator changes seem to still keep the overall signaling pathway activity at the levels observed in WT mice.

In *LRRK2* KO cortices, for instance, Wnt3a, pLRP6 (S1490) and BDNF protein levels were all significantly downregulated. This observation would be more in line with cortical *downregulation* of canonical Wnt signaling. In the cortex of G2019S KI mice, protein levels for Wnt3a and phosphorylated LRP6 (S1490)/LRP6 were elevated, while total LRP6, pGSK-3β (Y216) and BDNF were decreased. Phosphorylation of tyrosine 216 in GSK-3β has previously been shown to suppress Wnt activity (83). Therefore, most but not all of these changes point towards an expression profile more in line with *upregulated* canonical Wnt signaling activity in the cortex, which is in line with the overall *in vivo* findings.

Hippocampal samples revealed similar protein changes in *LRRK2* KO and G2019S KI samples, with a significant increase in β-catenin and a significant decrease in TCF1. The active β-catenin fraction was significantly reduced in KO hippocampi. In the striatum of *LRRK2* KO, mice gene expression for *β-catenin*, *Axin2*, and *Nfat* was significantly enhanced and protein levels of Wnt5a and LEF1 were significantly reduced. In KI striatum samples, protein levels of LRP6, phosphorylated GSK-3β (pY216)/GSK-3β, β-catenin, LEF1, and NFAT were significantly decreased, while the protein level of GSK-3β was augmented. Overall, these patterns suggest canonical Wnt signaling downregulation that was only confirmed by the significant change in active β-catenin/total β-catenin is the LRRK2 KO hippocampus.

In the for PD pivotal striatum, mRNA levels of signaling components in the PD LRRK2 G2019S KI mutant show no significant differences. This might be due to the opposing pattern of expression up- and downregulation observed in male and female mice. This pattern is diminished on protein level providing highly significant results, indicating a decrease in total Lrp6, GSK-3β (pY216)/GSK-3β, β-catenin, LEF1 and NFAT while showing an increase in total GSK-3β expression. This pattern speaks overall for conditions supporting the downregulation of canonical Wnt signaling in the striatum albeit without reaching any significant changes in active β-catenin/total β-catenin levels at 6-7 months of age.

In summary, whilst *in vivo* data as well as immunoblotted active β-catenin fractions suggest that *LRRK2* KO or pathogenic mutations may enhance Wnt activity, the finer molecular outlook underlying these phenotypes appears inconsistent across brain regions or *LRRK2* genotypes. There are two reasons which might account for the apparent discrepancy. Firstly, overall Wnt signaling activity tends to decline with aging (84). Thus, our *in vivo* observations may account for Wnt enhancements earlier in life which become less pronounced later on. It is true that our analysis revealed no individual effect of time. However, the molecular profiling experiments represent a single timepoint at 7 months of age, thus not taking into account earlier changes. Secondly, the *in vivo* data represents whole-brain activity, which encompasses a very complex microenvironment with intact cells of multiple types actively signaling. This scenario is not exactly reproduced in protein lysates from dissected brain areas. Therefore, it is possible that the overall *in vivo* observation of enhanced Wnt signaling might represent the summative activity of several positive and negative-feedback loops in different brain regions producing a net effect. This is an important challenge to the integration of *in vivo* and *ex vivo* data, which must be overcome in future studies.

One approach to clarify the significance, if any, of these discrepancies, should be to assess *in vivo* Wnt signaling *activation,* rather than basal levels alone, as was done here. Basal level analysis, whilst providing less physiological interference, is somewhat limited by relatively poorer signal-to-noise ratios. To initially address this, we employed another vector biosensor system *ex vivo,* in primary cultured cells (Figure 10).

Surprisingly, we did not observe any difference in basal Wnt signaling or NFAT signaling levels in cultured cortical neurons between WT, *LRRK2* KO or G2019S mice. However, whilst basal Wnt signaling levels were also comparable in cultured hippocampal neurons across all genotypes, basal NFAT signaling was enhanced in both mutant genotypes. In cortical neurons, Wnt activation via Wnt3a was actually dampened by *LRRK2* KO or G2019S, which is in contrast with our *in vivo* data. On the other hand, Wnt3a-mediated activation was enhanced by the G2019S mutation in hippocampal neurons, in line with whole-brain *in vivo* results. Taken together, these data are somewhat difficult to interpret, but initially suggest that individual cell type-specific Wnt activity phenotypes might differ from whole-brain signaling. Interestingly, it appears that only hippocampal neurons display patterns consistent with the *in vivo* experiments. It may be argued that these cells have a more significant impact on *LRRK2-*mediated Wnt phenotypes than previously thought.

It is important to note that our mouse models showed no phenotypical changes and all experiments were conducted under basal conditions. Therefore, future efforts should seek to address these issues by combining the *in vivo* approach with stimulation experiments. For instance, mice of different *LRRK2* genotypes expressing the biosensors could be treated with Wnt activating compounds such as lithium or Wnt ligands and/or immune stimulation via LPS or INFγ, and imaged long-term using the IVIS system. The resulting data might further clarify the *in vivo* role of LRRK2 in Wnt modulation with increased resolution.

Further studies should also seek to answer these questions in older mice, to even better reproduce the molecular environment of an ageing brain. In this respect, we recommend future investigations of mice up to 18-24 months of age.

## Conclusions

In conclusion, we have developed a novel method to investigate *in vivo* Wnt and NFAT signaling activity in PD mouse models, combining lentiviral biosensors with luciferase assays. Our data shows the system works reliably, with constructs delivered selectively to multiple brain regions, and *in vivo* changes supported by protein-level analysis. Our results support previous knowledge of a key Wnt and NFAT-regulatory role for LRRK2. There remain some discrepancies between the pictures shown by overall Wnt activity patterns and finer molecular dissection of the cascades. These should be addresses in subsequent work. Beyond our present findings, our experimental system is in itself a step forward in establishing non-invasive protocols to assess cerebral signaling pathway function *in vivo.* The potential applications are countless, from further elucidating molecular pathology, as done here, to testing candidate therapeutic compounds. We hope future work might build upon this notion to further our understanding of PD.

## Supporting information

Primer information

## Abbreviations

ABC: Activated β-catenin
ANOVA: Analysis of variance
BCA: Bicinchoninic acid
BDC: β-catenin destruction complex
BDNF: Brain-derived neurotrophic factor
CaN: Calciuneurin
CCD: Charge-coupled device
CD: Crohn’s disease
cPPT: Central polypurine tract
DLGN: dorsal lateral geniculate nucleus
DVL: Disheveled
FBS: Fetal bovine serum
Fz: Frizzled
GAPDH: Glyceraldehyde 3-phosphate dehydrogenase
GFP: Green fluorescent protein
GSK-3β: Glycogen synthase kinase 3β
GTP: Guanosine triphosphate
HBSS: Hank’s buffered saline solution
HPRT: Hypoxanthine-guanine-phosphoribosyl transferase
IFN-γ: Interferon-γ
IL: Interleukin
IP3: Inositol 1,4,5-triphosphate
KI: Knock-in
KO: Knockout
LPS: Lipopolysaccharide
LRP5/6: Low-density lipoprotein receptor-related protein 5/6
LRRK2: Leucine-rich repeat kinase 2
LTR: Long terminal repeat
NFAT: Nuclear factor of activated T-cells
NRON: Non-coding RNA repressor of NFAT
PBS: Phosphate buffered saline
PCR: Polymerase chain reaction
PCP: Planar cell polarity
PD: Parkinson’s disease
PLC: Phospholipase C
PVDF: Polyvinylidine fluoride
RE: Response element
ROI: Region of interest
SEM: Standard error of the mean
SFFV: Spleen focus-forming virus
TBS: Tris-buffered saline
TCF/LEF: T-cell specific transcription factor/lymphoid enhancer binding factor
Wnt: Wingless/Integrated
WPRE: Woodchuck hepatitis virus post-transcriptional regulatory element
WT: Wild type.

## Declarations

### Ethics approval and consent to participate

All animal investigations were approved by the University College London Animal Welfare and Ethical Review Body and licensed by the UK Home Office (PPL 80/2486 and PPL 70/9070).

### Consent for publication

Not applicable

### Availability of data and materials

Raw data is available upon request.

### Authors’ contributions

AW, AAR and KH designed the experiments; AW, SHL, TL, MH and YP performed the experiments; AW, KH, SHL and SG analyzed the data; TM and SW provided the lentiviral constructs; AW, SHL and SG made the figures; AW, KH, SHL and SG contributed to writing the submitted manuscript draft. All authors were involved in revising the manuscript and gave final approval of the version to be published.

## Acknowledgements

We are thankful to Dr Giulia Massaro for introducing the GFP staining technique. We further thank Professor Li Wei for constructive and helpful discussions about statistical testing. We acknowledge our colleagues from the UCL School of Pharmacy animal facility for their technical support and assistance in taking such good care for our mice.

---

**Figure S1.**
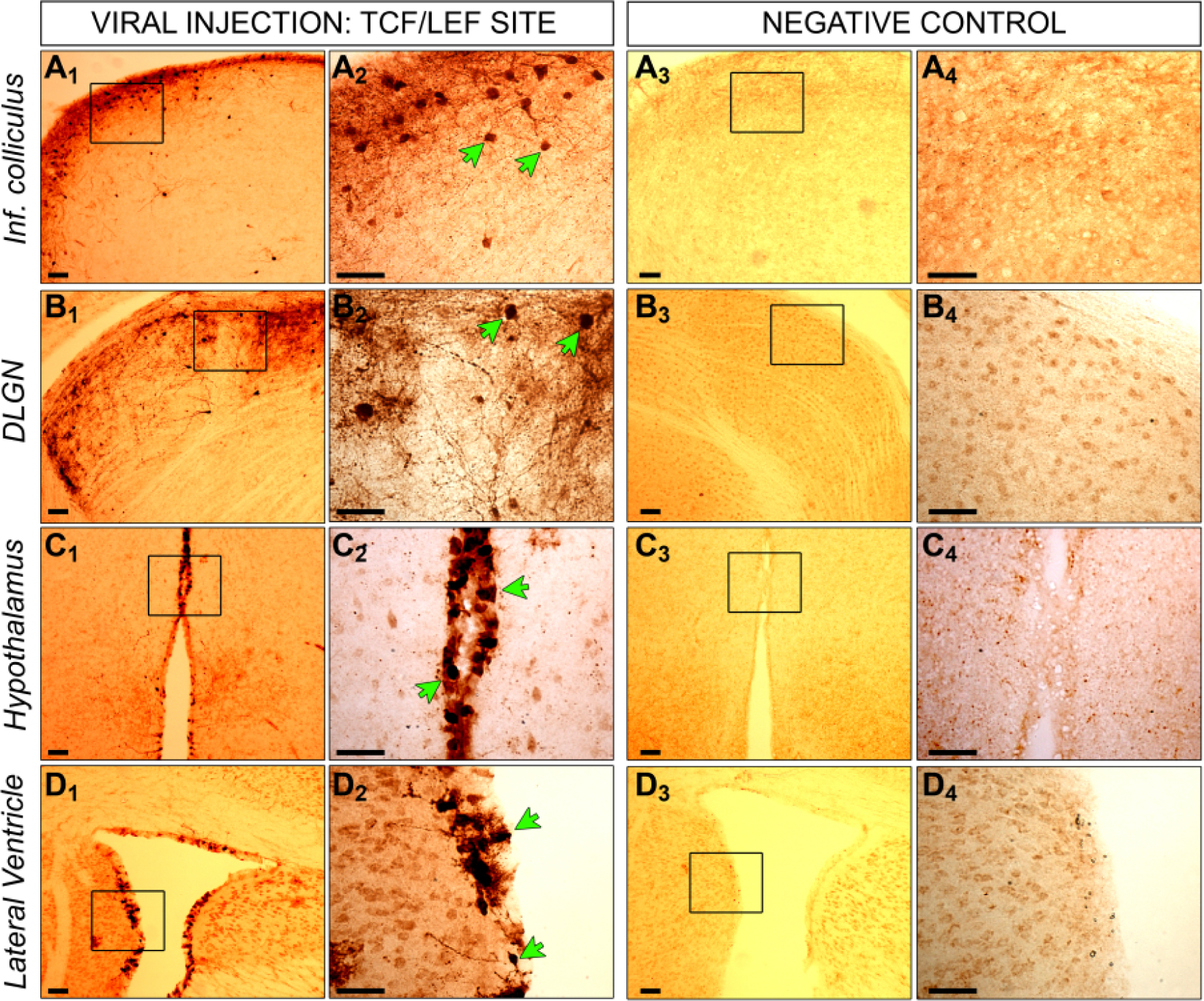
GFP staining of injected and non-injected mice with the TCF/LEF biosensor. Mice were injected intracranially at P0 with a lentiviral construct, containing a RE site for TCF/LEF and a second gene for GFP. Six months later, brains were collected from injected (**1** and **2**) and non-injected control mice (**3** and **4**). Brains were fixed in PFA, coronally cryosectioned into 40µm thick slices, which were stained via immunohistochemistry for GFP. Positive cells for GFP expression are visible in dark brown indicated by the green arrows. Positive cells were detectable in inferior colliculus (**A**), dorsal lateral geniculate nucleus (**B**), hypothalamus (**C**) and lateral ventricle (**D**). No positive cells were measurable in corresponding regions of non-injected control brains. Images in **2** and **4** represent the zoom of indicated squares in **1** and **3**. Scale bar for all images is 50µm.

**Figure S2.**
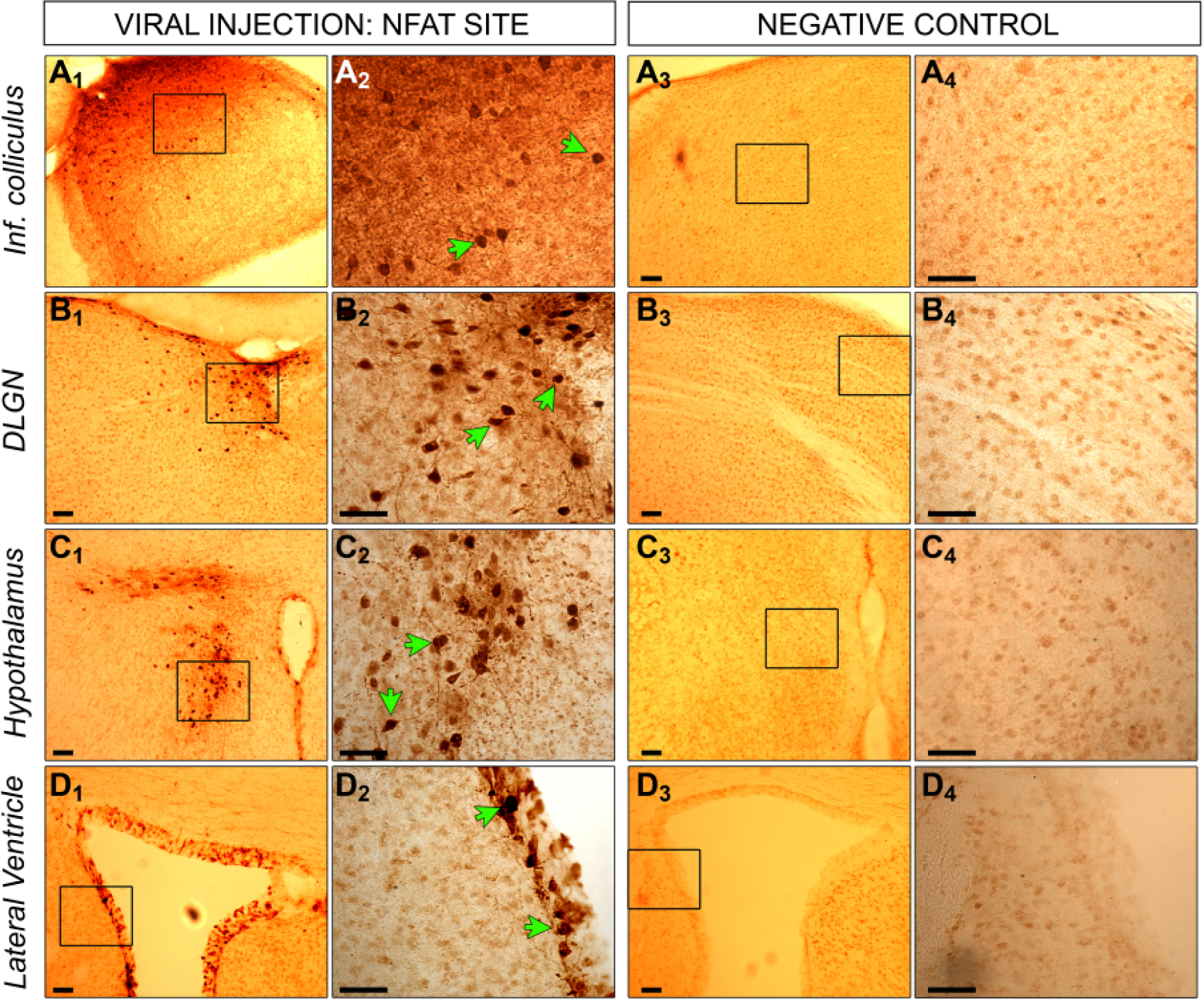
GFP staining of injected and non-injected mice with the NFATc1 biosensor. Mice were injected intracranially at P0 with a lentiviral construct, containing a RE site for NFATc1 and a second gene for GFP. Six months later, brains were collected from injected (**1** and **2**) and non-injected control mice (**3** and **4**). Brains were fixed in PFA, coronally cryosectioned into 40µm thick slices, which were stained via immunohistochemistry for GFP. Positive cells for GFP expression are visible in dark brown indicated by the green arrows. Positive cells were detectable in inferior colliculus (**A**), dorsal lateral geniculate nucleus (**B**), hypothalamus (**C**) and lateral ventricle (**D**). No positive cells were measurable in corresponding regions of non-injected control brains. Images in **2** and **4** represent the zoom of indicated squares in **1** and **3**. Scale bar for all images is 50µm.

