## Supplementary material for "Dysregulated Wnt and NFAT signaling in a Parkinson’s disease LRRK2 G2019S knock-in model": Primer information

| **Target Gene** | **Accession Number** | **Primer Sequence (5’-3’)** | **Amplicon** |
| --- | --- | --- | --- |
| *Wnt5a* | NM_009524.4 | (F) GTGCCATGTCTTCCAAGTTC  (R) TGCCTGTCTTCGCACCTT | cDNA: 246bp |
| *Wnt7a* | NM_009527.3 | (F) CGCCAAGGTCTTCGTG  (R) AGCCTAAGCTCTCGGAACTGT | cDNA: 191bp |
| β*-catenin* | NM_007614.3 | (F) CATTGGTGCCCAGGGAGA  (R) GATCAGGCAGCCCATCAACT | cDNA: 170bp |
| *Axin2* | NM_015732.4 | (F) GCCGACCTCAAGTGCAAACT  (R) GCCGGAACCTACGTGATAA | cDNA: 156bp |
| *Gsk-3* | NM_019827.7 | (F) TTGGACAAAGGTCTTCCGGC  (R) AAGAGTGCAGGTGTGTCTCG | cDNA: 349bp |
| *Tcf1* | NM_001313981.1 | (F) ATCTGCTCATGCCCTACCCA  (R) GCGGCCTGTGAACTCCT | cDNA: 192bp |
| *Nfatc1* | NM_016791.4 | (F) CGGGAAGAAGATGGTGCT  (R) CTGGTTGCGGAAAGGTGGTA | cDNA: 169bp |
| *Bdnf* | NM_001048139.1 | (F) CGGGACGGTCACAGTC  (R) CCGAACATACGATTGGGTAG | cDNA: 166bp |
| *Gapdh* | NM_008084.3 | (F) GCCCAGAACATCATCCCT  (R) GTCCTCAGTGTAGCCCAAGA | cDNA: 231bp  gDNA: 315bp |
| *Hprt* | NM_013556.2 | (F) AGTCCCAGCGTCGTGATTAG  (R) GGGCCACAATGTGATG | cDNA: 181bp |
| β*-actin* (51) | NM_007393.5 | (F) GCCAACCGTGAAAAGATGAC  (R) GGCGTGAGGGAGAGCATAG | cDNA: 183bp  gDNA: 637bp |
| *Titin* | NM_011652.3 | (F) TGCTGGAACTCCCGAGATTG  (R) CTCCGCTGTCGCTGCTATCA | gDNA: 137bp |
| *WPRE* | JC285214 | (F) TTGCTTCCCGTATGGCTTTC  (R) GCCGTGGCAATAGGGAGG | vDNA: 202bp |

**Table 1 Primer information for quantitative real time polymerase chain reaction**
